# Structural Characterization of Calcium-Dependent Calmodulin-Calmidazolium Binding using Capillary Vibrating Sharp-Edge Spray-based Native Mass Spectrometry and In-Droplet Hydrogen Deuterium Exchange Mass Spectrometry

**DOI:** 10.64898/2026.05.15.725515

**Authors:** Sohag Ahmed, Axel Woehrling, Peng Li, Stephen J. Valentine, Kevin C. Courtney

## Abstract

Capillary vibrating sharp-edge spray ionization (cVSSI)-based native mass spectrometry (nMS) and in-droplet hydrogen deuterium exchange (HDX-MS) were used to evaluate calcium-dependent interactions between calmodulin and calmidazolium (CDZ). cVSSI was found to produce a narrow charge-state distribution (CSD) with low average charge states, indicating that this method preserved native-like states. cVSSI was also able to resolve stepwise Ca^2+^ binding for species containing one to four Ca^2+^ ions. In the absence of Ca^2+^, no detectable CDZ binding was observed. However, CDZ binding was observed when calmodulin was fully loaded with Ca^2+^. CDZ binding to the protein caused marked redistribution of the CSD toward lower charge states, consistent with ligand-induced stabilization of the protein into a more compact conformational ensemble. The apparent dissociation constant (*K_d_*) of the interaction was determined to be 261 ± 29 nM and 126 ± 17 nM from Langmuir and quadratic binding models, respectively. Complementary in-droplet HDX-MS revealed that deuterium uptake was correlated with charge state across the CSD, corroborating the nMS evidence for stepwise conformational compaction of CaM. Within the individual charge states, two distinct populations representing CDZ-bound and unbound CaM were resolved by HDX-MS, which were not detectable by nMS alone. Comparison of these populations within each of the 6+, 5+ and 4+ charge states revealed reductions in deuterium uptake by approximately 37%, 22%, and 11% upon CDZ binding, respectively, consistent with ligand occlusion and local structural tightening at the binding interface. The gradient in protection across charge states further reflects differences in binding interface accessibility among CaM conformers spanning a range of CDZ-induced conformational states. Together, these results demonstrate that cVSSI-based nMS coupled with in-droplet HDX-MS provides an integrated platform for simultaneously resolving metal loading, ligand binding, binding affinity, and ligand-induced conformational changes. This approach complements traditional structural methods by enabling direct interrogation of dynamic, metal-dependent protein–ligand interactions in their native states.

## INTRODUCTION

Native mass spectrometry (nMS) has become a powerful analytical platform to characterize intact biomolecules and their complexes under near-physiological conditions.^1–8^ The development of soft ionization tools, especially electrospray ionization (ESI), has enabled the transfer of large biomolecules from solution to the gas phase while retaining the molecular integrity and, in some instances, non-covalent interactions between two or more molecules.^7,9–12^ This feature has made nMS a valuable tool to study proteins, nucleic acids, and their complexes under conditions favoring and approaching their native states; such studies conducted under even mild denaturing conditions in which conformer ensembles shift toward less native states are not accessible with other structure determination techniques.^7,13–15^ Therefore, nMS is a widely used approach to investigate biomolecule conformer ensembles and their interactions as it can provide structural insight together with information on binding stoichiometry, metal-ion occupancy, oligomeric form, and the presence of co-existing conformational populations in a single experiment.^16–21^

Although nMS is widely used for structural characterization of biomolecules, the efficiency of the method depends strongly on the ionization technique employed.^22–25^ Furthermore, the preservation of the native state of biomolecules from solution to the gas phase depends on the ionization source utilized as well as the parameters under which the source is operated.^26–28^ Nanoelectrospray ionization (nESI)^29^ is the most commonly used ion source in nMS because of its low flow rate, improved sensitivity, and relative compatibility with volatile aqueous buffer solutions.^30–32^ However, conventional ESI and nESI can present limitations, including salt adduction^33^, source-induced activation^34,35^, and, in some cases, distortion of the conformational ensemble during ion production and transmission into the mass spectrometer^27,36^. These limitations are more critical for fragile or metal-dependent systems, where retention of the native state and weak non-covalent interactions are essential for structural studies of biomolecules.^19,37^ Therefore, continued development of a gentler and more tunable ionization method remains an important goal in nMS studies.

Recently, spray-based ionization sources that do not employ an electric field, including surface acoustic wave ionization^38,39^ and mechanospray ionization (MOSI)^40^, have shown a remarkable ability to preserve the native structure of proteins. Capillary vibrating sharp-edge spray ionization (cVSSI) is another such alternative that may offer several advantages over conventional methods such as ESI/nESI.^27,36,41^ In cVSSI, droplet production is driven mechanically rather than solely through the high-voltage-driven Taylor cone mechanism used in ESI/nESI, allowing spray generation with little or no applied electric field and providing greater flexibility in controlling ionization behavior.^36,41^ Previous studies demonstrated that cVSSI could preserve fragile protein structures better than conventional voltage-driven ESI/nESI ionization, producing lower average charge states and narrower charge-state distributions for globular proteins under near native conditions.^27,36,42^ These findings established cVSSI as a soft ionization source for nMS and suggested that it may better retain native-like biomolecular structure during transfer to the gas phase. More recently, voltage-tunable cVSSI was shown to control droplet charging and thereby alter the conformer ensembles sampled by MS; this revealed that low-, mid-, and high-voltage conditions can favor distinct protein populations and charge-state distributions.^27,36,41^ Together, these studies indicate that cVSSI is not only a gentle ionization source but also a tunable platform for studying how ionization conditions influence conformer preservation.

In negative ion mode, applying an electric field during cVSSI was shown to improve ion production relative to conventional ESI, particularly for aqueous samples where higher voltages used in ESI can trigger corona discharge and compromise signal stability and sensitivity.^43^ This improvement is relevant to native-state studies of biomolecules whose functional states depend on the presence of salt or metal ions in solution.^41,44–46^ For example, recent work on triplex DNA demonstrated that cVSSI can be optimized to enhance native-like ion production while reducing ion adduct formation and preserving higher-order structure.^41^ Furthermore, this previous study also showed that control of source voltage and inlet capillary temperature can significantly improve the quality of nMS measurements.^41^ Thus, the prior work has established cVSSI as a useful platform for the study of peptides^47^, proteins, protein conformer ensembles^48^, and structured nucleic acids^41^ under near-physiological buffer conditions.

A complementary advancement in MS studies was the development of cVSSI-enabled in-droplet hydrogen-deuterium exchange MS (HDX-MS).^42,48^ In conventional HDX-MS experiments, analyte solutions are pre-incubated with D_2_O for the exchange reaction before spraying to the MS inlet.^49–53^ HDX-MS has been widely used to probe protein dynamics and ligand-induced conformational change by monitoring the hydrogen-deuterium exchange behavior.^53–56^ Inspired by rapid HDX-MS experiments employing theta capillaries,^57,58^ the cVSSI-enabled approach uses separate plumes of analyte and D_2_O that are generated with two cVSSI devices; this allows the sample to mix immediately prior to MS sampling, enables rapid exchange within the microdroplets, improves temporal resolution and enhances detection of transient conformational states.^42,48,59–63^ Earlier cVSSI-based in-droplet HDX studies showed that the approach can distinguish peptide structures with different helical configurations, reveal coexisting conformers of ubiquitin in mildly denaturing to denaturing solutions, and provide information on protein structural heterogeneity.^48^ In addition, this approach was extended to DNA systems, demonstrating that in-droplet HDX could differentiate folded and unfolded nucleic acid conformers and probe relative protection among quadruplex, duplex, and triplex DNA structures.^64^ More recently, in-droplet HDX combined with molecular dynamics simulations was used to relate rapid HDX behavior to peptide conformational flexibility and structural bias, supporting the broader notion that the method can serve as a rapid structural probe for biomolecular systems.^47^ Collectively, these studies suggest that cVSSI-based in-droplet HDX-MS can complement nMS by providing an additional readout of structural flexibility, protection, and binding-induced conformational changes.

In this study, cVSSI-based nMS and in-droplet HDX-MS have been extended to a protein-ligand system in which ligand binding depends on the metal occupancy and conformational state of the protein. Calmodulin (CaM) is an ideal model for this type of study as it is a relatively small and well-characterized protein. CaM is a eukaryotic calcium-binding protein that functions as a central sensor of intracellular Ca^2+^ signaling.^65^ It contains four canonical EF-hand Ca^2+^ binding motifs arranged as two lobes connected by a flexible linker.^66^ Calcium binding induces conformational rearrangement, exposing hydrophobic surfaces and enabling interactions with a wide range of proteins and small molecules.^66,67^ As the structural and functional state of CaM is tightly coupled with Ca^2+^ loading, and ligand binding causes significant structural rearrangement, CaM is used herein as a prototypical metal-dependent binding system to evaluate the utility of cVSSI-based nMS and in-droplet HDX-MS methods.

Calmodulin is a highly promiscuous Ca²⁺ sensor that interacts with hundreds of target proteins and is known to bind a wide range of small-molecule inhibitors.^68^ Calmidazolium (CDZ) is one of the most well-known CaM inhibitors and preferentially binds to the Ca^2+^-loaded form of the protein.^69^ This system is well-suited for evaluating protein–ligand interactions that are coupled to metal loading using cVSSI-based nMS and in-droplet HDX-MS. In particular, the cVSSI-based nMS can resolve stepwise metal binding, distinguish ligand-bound and unbound populations, and provide an apparent dissociation constant (*K_d_*) from titration-based measurements of bound and unbound protein species. Furthermore, in-droplet HDX-MS can reveal ligand-induced structural changes in solvent protection and dynamics, and has the potential to resolve conformationally distinct populations that are indistinguishable by nMS alone.

In the present work, cVSSI-based nMS was applied to characterize CaM under near native conditions, resolve stepwise Ca^2+^ loading and evaluate the Ca^2+^-dependent interaction of CDZ with CaM. Using the CaM-Ca^2+^-CDZ system, nMS identified CaM-Ca^2+^ binding and showed that CDZ predominantly bound CaM when it was fully loaded with Ca^2+^, revealing a clear dependence of ligand binding on metal occupancy. Titration experiments enabled estimation of the apparent dissociation constant (*K_d_*) of the interaction from the relative abundances of bound and unbound species, which fell within the range of prior reports.^69–71^ Additionally, in-droplet HDX-MS corroborated the nMS evidence for stepwise conformational compaction of CaM and was further able to resolve distinct CDZ-bound and unbound populations within individual charge states, revealing charge-state-dependent differences in ligand-induced protection consistent with CDZ binding across a range of coexisting conformational states. Altogether, these results highlight the utility of cVSSI-based approaches for probing protein-ligand binding stoichiometry, apparent binding affinity, and conformational stabilization across heterogeneous protein ensembles.

## EXPERIMENTAL SECTION

### Chemical and Reagents

Ammonium acetate (ACS grade) was obtained from Fisher chemical (Waltham, MA, USA). Calcium acetate was purchased from Thermo Fisher Scientific (Ward Hill, MA, USA). Deuterium oxide (D_2_O) was purchased from Sigma-Aldrich (St. Louis). Dimethyl sulfoxide (DMSO) was purchased from Fisher Scientific. Calmidazolium chloride (CDZ) was purchased from MedChemExpress LLC (Monmouth Junction, NJ, USA). All chemicals were used without any further purification. The protein was expressed and purified in the laboratory. Spectra/Por molecular porous membrane tubing (6-8 kDa) was obtained from Spectrum Laboratories, Inc (Rancho Dominguez, CA, USA).

### Protein expression and purification

The human CaM (hCaM) cDNA was acquired from IDT as a custom gBlock synthesis and cloned into a PCR-linearized pET-SUMO vector using the In-Fusion enzyme cloning master mix from Takara. Recombinant hCaM was expressed and purified from *E. coli* by following a previously published protocol from our group.^72^ Briefly, the pET-SUMO-hCaM vector was transformed into BL21 cells and propagated in 50mL of LB media overnight. This 50mL starter culture was then split evenly between six Fernbach flasks containing 1L LB and grown at 37 °C until the media reached an OD_600_ of 0.6-0.8. SUMO-hCaM expression was then induced by 200 mM IPTG and incubated for another 3 hours at 37 °C, followed by centrifugation to harvest the E. coli pellet. The cells were then lysed by sonication, and the cell debris was pelleted by centrifugation to isolate the soluble protein fraction in the supernatant. Histagged SUMO-hCaM was then affinity-purified using Hispur cobalt resin and cleaved from the resin by hSENP2 to liberate the hCaM from the SUMO domain. The isolated protein was then further purified by FPLC using a Superdex 75 increase 10/300 column and quantified by SDS-PAGE. Prior to mass spectrometry analysis, the hCaM was dialyzed into 100 mM ammonium acetate.

### Sample Preparation

A stock of 126 µM CaM protein was prepared and diluted to the required concentration (0.5 µM) from the stock. Ammonium acetate was dissolved in Mili Q water to prepare a 100 mM ammonium acetate solution. Originally, the protein was in resuspension buffer (50 mM TRIS, 300 mM NaCl, 10% glycerol, pH 7.4), which was exchanged to 100 mM ammonium acetate using Spectra/Por molecular porous membrane tubing (6-8 kDa MWCO). The solution was replaced every 4 hours (×3) to completely exchange the buffer to ammonium acetate. A stock of 10 mM CDZ was prepared by dissolving it in DMSO. The stock solution was diluted based on the requirements of the experiments. DMSO content in the sample was maintained at 0.2% (v/v) consistently by adding 1 µL of the CDZ solution to 500 µL of protein solution for each of the ligand addition experiments.

### cVSSI device Fabrication for nMS and HDX

The cVSSI devices were fabricated following the previously described procedures for nMS and in-droplet HDX-MS experiments.^41^ For this study, microdroplet plumes were generated at an RF frequency of 92-96 kHz and post-amplification amplitude of 8.5-8.8 Vpp. The flow rate of the syringe pump was maintained at 10 µL/min. For field-enabled cVSSI experiments in positive-ion mode, a DC bias of 1300 V was applied to the sample solution through the platinum wire inserted into the poly-tetrafluoroethylene tube. Approximately 2-3 mm distance was maintained between the inlet capillary and the cVSSI device tip.

For HDX-MS experiments, two devices were employed to generate separate plumes for the analyte and D_2_O solutions (Figure S1). Each solution was infused at a rate of 10 µL/min. The RF frequency and amplitude were maintained the same as those used in the nMS experiments. The dual-emitter and overall experimental configuration were identical to the previously reported setup.^36,73^

### MS Settings

All the nMS and in-droplet HDX-MS experiments were performed on a Q-Exactive hybrid quadrupole-orbitrap mass spectrometer (Thermo Fisher Scientific, San Jose, CA). The data were acquired in positive ion mode with a scan range of 1000-6000 *m*/*z* and the resolving power was set to 140,000. The inlet capillary temperature was set at 300°C, while the maximum injection time and automatic gain control (AGC) target were set at 10 milliseconds and 1×10^6^, respectively.

### Data Analysis

Mass spectra data were analyzed using the XCalibur Qual Browser software (Thermo Scientific). To generate mass spectra, each of the datasets were signal averaged and the x,y values were extracted and imported into Excel worksheets for further analysis and plotting.

For the determination of the apparent dissociation constant (*K_d_*), each of the datasets in different concentrations was processed in Excel, and the peak intensities of the ligand-bound, and unbound peaks were recorded and fraction of protein bound was determined using Equation (1).^74^

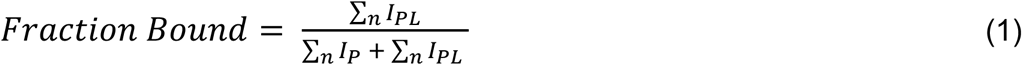

Here, ∑*n I_P_* and ∑*n I_PL_* represent the sum of the intensity of the unbound protein ions and the sum of the intensity of the ligand-bound protein ions across the resolved charge states (where n = 4+ and 5+), respectively.

To determine the apparent *K_d_*, two independent methods were used: the Langmuir binding model (Equation 2)^75^ and the quadratic binding model (Equation 3)^76^.

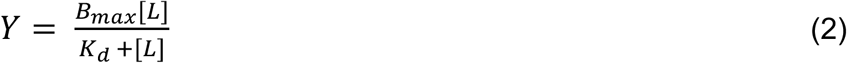

In Equation 2, *Y* = Predicted values, *B*_max_ = Maximum fraction bound, [*L*]= Ligand concentration, and *K_d_* = Apparent equilibrium dissociation constant.

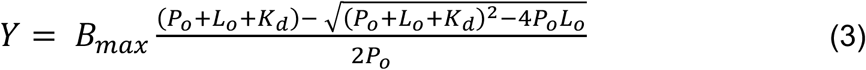

In Equation 3, *Y*= Predicted values from the quadratic binding model, *B_max_*= Maximum fraction bound, *P_o_*= concentration of the protein, *L_o_*= initial concentration of the ligand and *K_d_*= apparent equilibrium dissociation constant of the interaction.

The number of incorporated deuterium atoms was determined from the shift of the weighted average m/z values after HDX exposure and multiplying the shift by the respective charge state. The weighted average m/z of a single charge state was calculated using Equation (4)

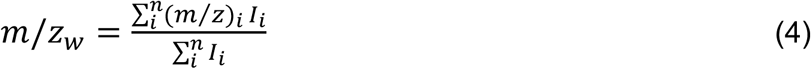

Here, (*m*⁄z)*_i_* and *I_i_* represent the mass-to-charge ratio and intensity of the *i*th isotopic peak, respectively, and *n* is the total number of isotopic peaks considered in the distribution.

### Structural Modeling and Analysis

Predicted protein structures were made using Chai-1 via the Chai Discovery web interface^77^ (https://lab.chaidiscovery.com/dashboard) and default parameters were used. Five structures were modeled: (I) apo CaM, (II) CaM and four Ca^2+^ ions, (III) CaM and five Ca^2+^ ions, (IV) CaM and four Ca^2+^ ions and one CDZ molecule, (V) CaM and four Ca^2+^ ions and two CDZ molecules. Five structural models were generated for each of these five structures.

The UniProt amino acid sequence P0DP23 · CALM1_HUMAN was inputted to the Chai-1 interface to generate the human CaM protein, and was used without modification unless otherwise specified. The “protein” option was chosen for the “molecule type” parameter when entering the CaM amino acid sequence. The SMILES sequences [Ca+2] and ClC1=CC=C(C=C1)[C@@H](C1=CC=C(Cl)C=C1)N1C=CN=C1C[C@@H](C1=CC=C(Cl)C=C1)C1=CC=C(Cl)C=C1 were used to generate Ca^2+^ ions and CDZ molecules, respectively. The “ligand” option was chosen for the “molecule type” parameter when entering both the Ca^2+^ ion and CDZ molecule sequences.

As shown in Table S1, the predicted structures were compared to experimentally determined crystal structures obtained from the Protein Data Bank (PDB). The CaM + 4 Ca^2+^ AI-predicted structure was compared to the PDB ID entry 1CLL^78^, the CaM + 1 CDZ + 4 Ca^2+^ AI-predicted structure was compared to the PDB ID entry 7PSZ^69^, and the CaM + 2 CDZ + 4 Ca^2+^ AI-predicted structure was compared to the PDB ID entry 7PU9^69^.

The UCSF ChimeraX^79^ software was used to align and analyze the structures. The PDB crystal structures were missing residues as follows: both 1CLL and 7PU9 lack residues 1-4 (MADQ) and 149 (K) and 7PSZ lacks residues 1-2 (MA) and 148-149 (AK). These residues were deleted from the corresponding predicted models before alignment to have equivalent alignment coverage and to avoid Root-mean-square deviation (RMSD) inflation. Specifically, residues 1-4 (MADQ) and 149 (K) were deleted before alignment for the CaM + 4 Ca^2+^ and CaM + 2 CDZ + 4 Ca^2+^ predicted models, and residues 1-2 (MA) and 148-149 (AK) were deleted before alignment for the CaM + 1 CDZ + 4 Ca^2+^ models.

The ChimeraX MatchMaker tool was used to perform structural superposition and for analysis. The following parameters were used: bb chain pairing, Needleman–Wunsch alignment algorithm, BLOSUM-62 similarity matrix, 0.3 secondary structure fraction, 18/18/6 gap open (HH/SS/other), 1 gap extend, ((6-9-6), (6-6), (4)) (HSOxHSO) secondary structure matrix, and 2 Å iteration cutoff. Each predicted model was superposed and compared to its corresponding crystal structure. For each comparison, both the RMSD value between pruned atom pairs (Pruned RMSD) and the RMSD value across all pairs were calculated. The model with the highest coverage ([pruned pairs / total pairs] × 100) relative to its reference crystal structure was selected for the figures.

## RESULTS AND DISCUSSION

### Characterization of CaM Structure Preservation and Analysis of Ca^2+^-dependent CDZ Binding

In nMS studies, the presence of low charge states and a narrow charge-state distribution (CSD) is generally associated with the preservation of compact, native-like biomolecule conformations. In contrast, broader and higher CSDs are associated with unfolded/denatured conformations.^80–84^ A comparative study between cVSSI and conventional ESI on globular proteins showed that cVSSI, under low-field conditions, can produce low charge states with a narrower CSD than ESI.^36^ Consistent with these observations, the mass spectrum of apo-CaM (Figure 1A) exhibits relatively low charge states and a narrow CSD, indicating preservation of the native-like state of the protein with cVSSI. The observed CSD for apo-CaM is primarily limited to charge states 6+ to 8+, suggesting preservation of the protein structure in the microdroplets and into the gas phase under native-like conditions. Previous nMS and IMS-MS studies of CaM reported a wider or multimodal CSD with low to higher charge states ( z = 7+ to 16+), where low charge states corresponded to compact/native-like conformers and higher charge states to unfolded conformers.^85–87^ Although previous ESI and nESI-based nMS studies identified apo-CaM as a dynamic protein with folded and unfolded conformational states, cVSSI-based nMS data from this study predominantly detects a narrow CSD with lower charge states reflecting native-like/folded conformational states. These findings suggest that cVSSI may better preserve the native-like state of the protein under these conditions, supporting its broad utility as a soft ionization technique.

**Figure 1.**
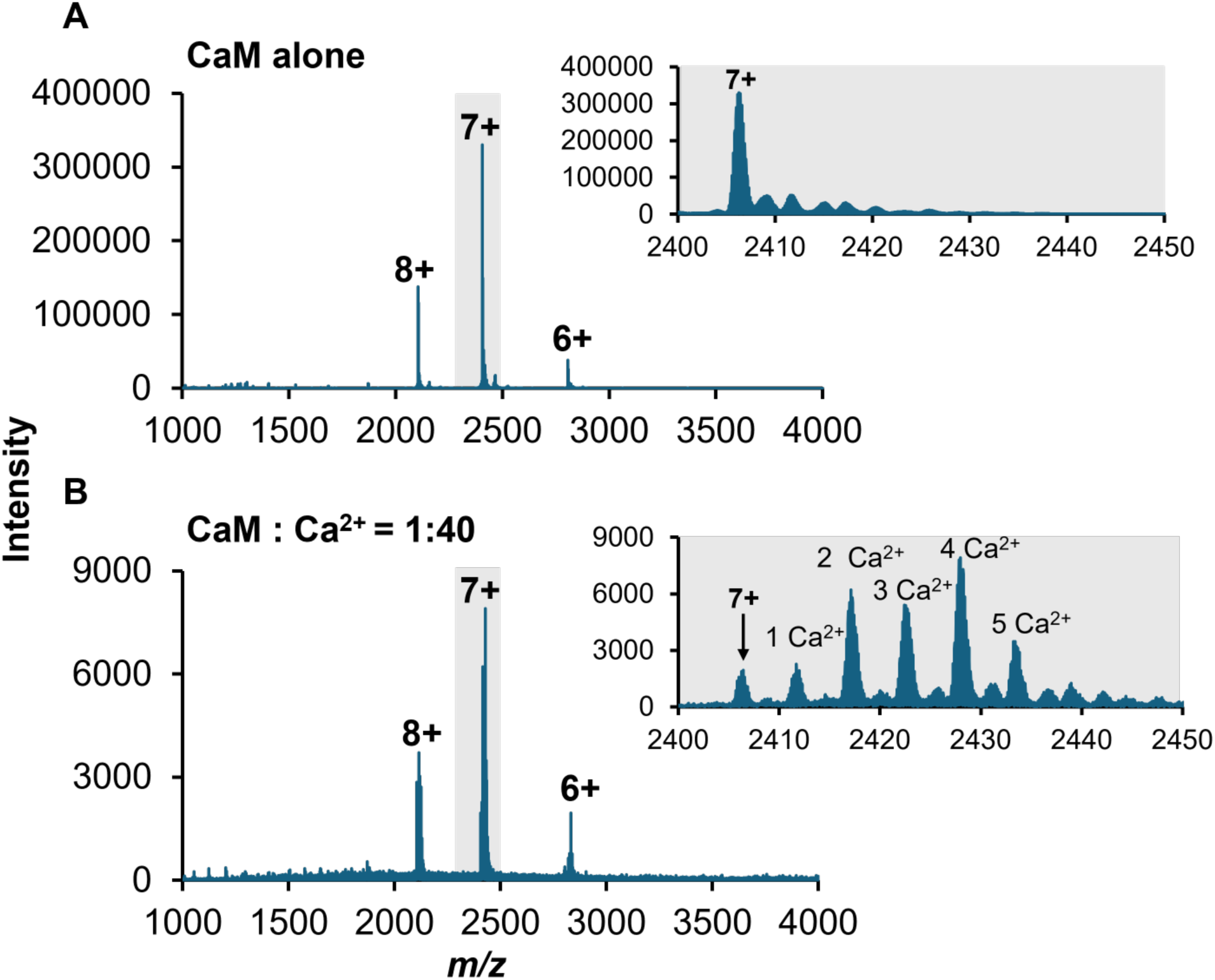
Native mass spectra of Calmodulin protein. The x-axis represents the mass to charge ratio (*m*/*z*) and the Y-axis the ion intensities. The left panel shows the mass spectrum of the protein in native state **(A)** vs Ca^2+^ bound state **(B).** Right panel shows the zoomed-in region of the 7+ charge state to better visualize the spectral difference and binding of the Ca^2+^ to the protein. The CaM and Ca(OAc)_2_ concentrations were maintained at 0.5 μM and 20 μM, respectively, throughout the experiment

Having confirmed the preservation of the native-like state of the protein, Ca^2+^ binding by CaM was next characterized using cVSSI. Calcium acetate (Ca(OAc)_2_) was added to the protein sample and incubated for 10 minutes before spraying into the MS inlet. Figure 1B presents the mass spectrum of Ca^2+^-bound CaM. In the spectrum, multiple new Ca^2+^-bound CaM species were observed, allowing resolution of Ca^2+^ binding stoichiometry by cVSSI-based nMS (Figure 1B). Distinct Ca^2+^-bound CaM species containing predominantly one to four Ca^2+^ ions were observed, suggesting stepwise occupation of the four canonical EF-hand calcium-binding sites of the protein (Figure 1). Previous nMS studies have also reported one to four Ca^2+^-bound species of CaM.^85^ Although, similar to a previous study by nESI/ESI^85,88^, fewer ions corresponding to CaM with five Ca^2+^ ions bound were also observed, we attribute this to limited non-specific binding as CaM contains only four Ca^2+^ binding sites.^89^ That said, structural predictions suggest that a fifth Ca^2+^ ion could, in principle, be transiently coordinated adjacent to one of the four canonical Ca^2+^-binding sites. However, the fifth Ca^2+^ was not localized to a consistent binding site across the models (Figure S2); it appeared in different regions of the protein, suggesting that the fifth Ca^2+^ ion may represent a non-specific association. Furthermore, considering CaM is well established to only bind four Ca^2+^ ions, this interaction is unlikely to be biologically relevant.

To determine the ratio of the protein to Ca(OAc)_2_ required to fully load the Ca^2+^ binding sites of the protein, different concentrations of Ca(OAc)_2_ were titrated against a fixed concentration (0.5 µM) of protein at CaM:Ca(OAc)_2_ ratios of 1:1, 1:4, 1:10, 1:20, and 1:40. When increasing the concentration of Ca(OAc)_2_, a progressive enrichment of the Ca^2+^ bound species was observed (Figure S3). At a ratio of 1:40 of the protein (0.5 µM) to Ca(OAc)_2_ (20 µM), the CaM species with four Ca^2+^ bound became the predominant feature, suggesting near saturation of the four Ca^2+^-binding sites of the protein under these experimental conditions. Having established that CaM became populated with a fully loaded Ca^2+^ form under these conditions, they were used to examine whether Ca^2+^ loading is required for CDZ binding.

The addition of CDZ to the sample in the absence of Ca^2+^ did not lead to the emergence of a new CaM:CDZ species in the mass spectrum. That is, no detectable binding to CaM was observed upon addition of CDZ without Ca^2+^ (Figure 2. A vs C). However, the mass spectrum (Figure 2C) suggested a moderate degree of conformer ensemble transition (perhaps compaction) upon the addition of CDZ solution, as the lower charge state 6+ species showed a higher intensity while the higher charge state 8+ ion intensity was comparatively reduced (see Figures 1 and 2). Furthermore, a minor 5+ species was observed for CaM upon addition of CDZ in the absence of Ca^2+^, which was previously undetected. Further analysis determined that these moderate structural modifications do not result from ligand interaction and are instead triggered by the CDZ solvent. The ligand was introduced into the protein from a DMSO stock (final DMSO concentration of 0.2% v/v). It has previously been demonstrated that a small quantity of organic solvent can affect the CSDs of proteins in mass spectra.^90,91^ Figure 3 demonstrates the direct effect of DMSO on the CSD of CaM in the presence and absence of the CDZ ligand. The change in the CSD observed upon CDZ addition without Ca^2+^ are similar to that observed with DMSO alone. These findings confirm the change in the CSDs of CaM with CDZ in the absence of Ca^2+^ does not result from transient ligand binding, but rather from the presence of a small quantity of DMSO. Thus, the data suggest that CDZ does not appreciably interact with CaM in the absence of Ca^2+^.

**Figure 2.**
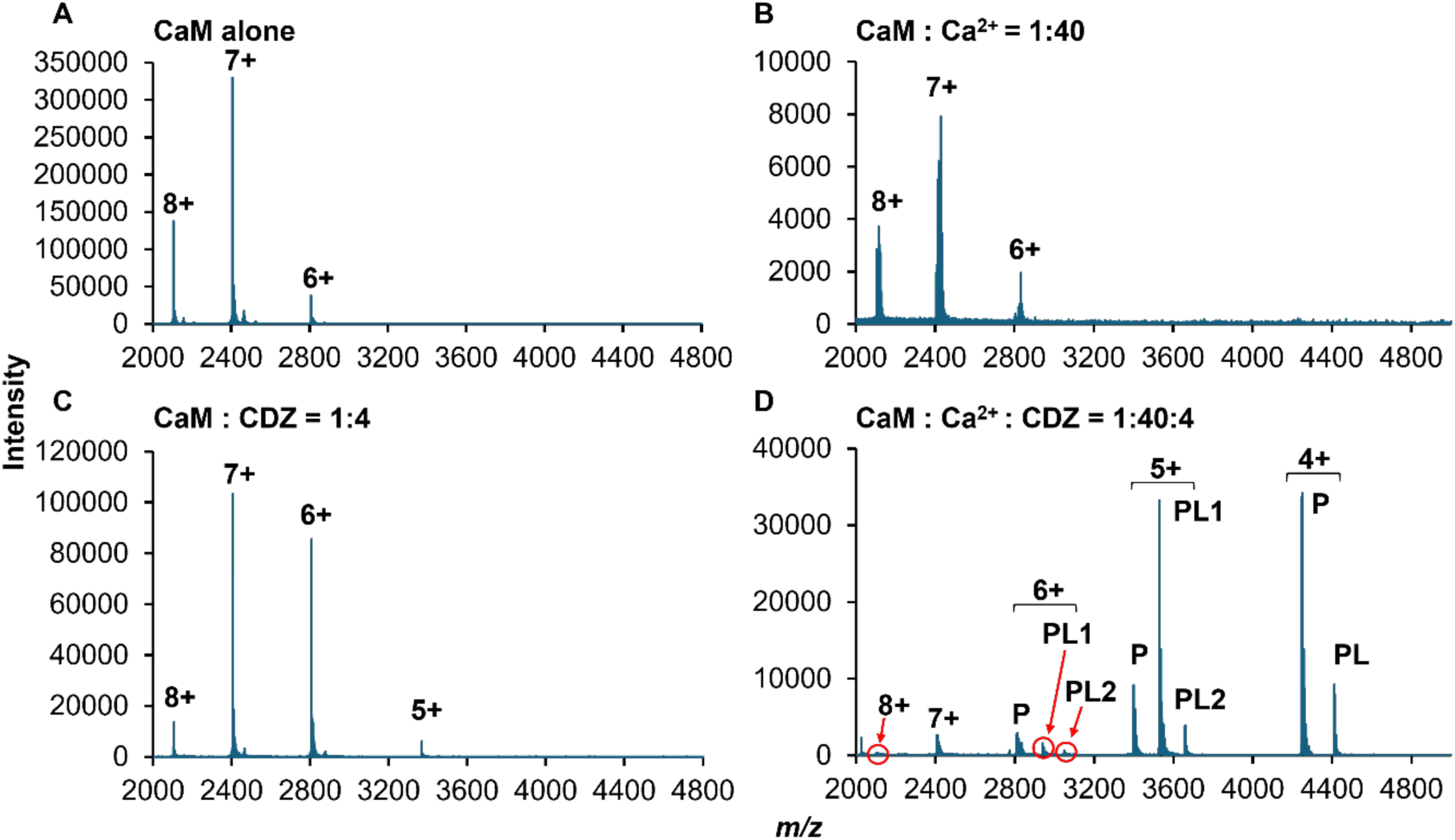
Native mass spectra of CaM acquired with and without CDZ in the absence and presence of Ca^2+^. The CaM, Ca(OAc)_2_ and CDZ concentrations were maintained at 0.5 μM, 20 μM, and 2 μM, respectively. **A.** CaM without CDZ in the absence of Ca^2+^. **B.** CaM without CDZ in the presence of Ca^2+^. **C.** CaM with CDZ in the absence of Ca^2+^. **D.** CaM with CDZ in the presence of Ca^2+^. P, PL1, and PL2 correspond to populations of CaM that are unbound, one and two CDZ molecules bound, respectively.

**Figure 3.**
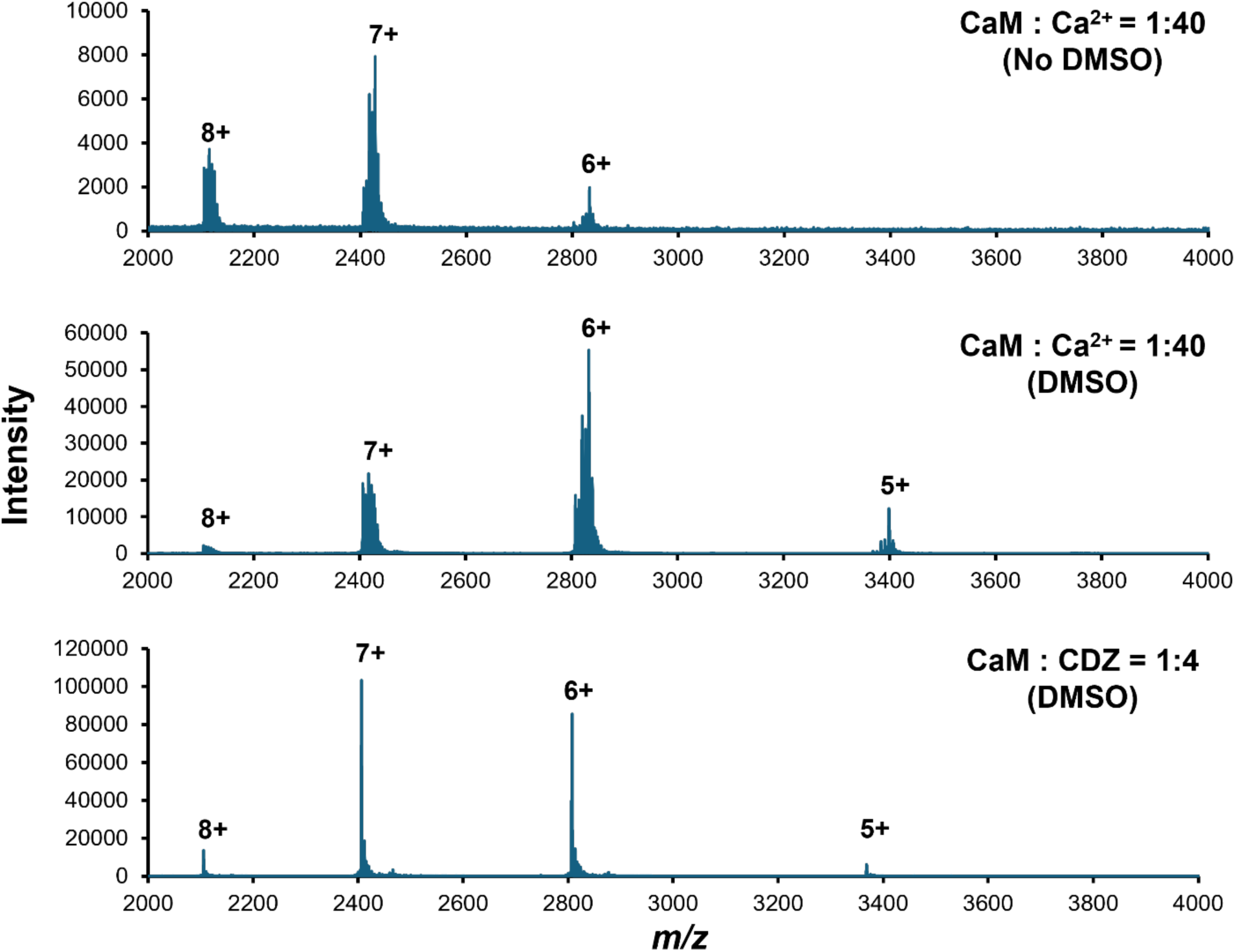
Effect of DMSO on the native mass spectra of CaM. The CaM, Ca(OAc)_2_ and CDZ concentrations were maintained at 0.5 µM, 20 µM, and 2 µM, respectively. Top to bottom: CaM with Ca^2+^ in the absence of DMSO (0.02 % v/v), CaM with Ca^2+^ in the presence of DMSO (0.02 % v/v), CaM with Ca^2+^ and CDZ in the presence of DMSO (0.02 % v/v). DMSO: Dimethyl Sulfoxide.

The samples containing CaM, CDZ, and Ca^2+^ were also analyzed. In contrast to the CDZ-free condition, which exhibited a heterogeneous Ca^2+^ binding stoichiometry, we found that the CDZ-bound complexes were predominantly fully loaded with Ca^2+^. After the addition of CDZ, CaM species with one to three bound Ca^2+^ ions were no longer detected, and only the fully Ca^2+^-loaded form remained (Figure 4). These findings suggest that CDZ preferentially binds to the fully Ca^2+^-loaded population and are consistent with ligand-induced conformational stabilization of the protein. This behavior further suggests thermodynamic coupling between Ca^2+^ binding and ligand association, whereby CDZ binding shifts the equilibrium toward the fully Ca^2+^-saturated state.

**Figure 4.**
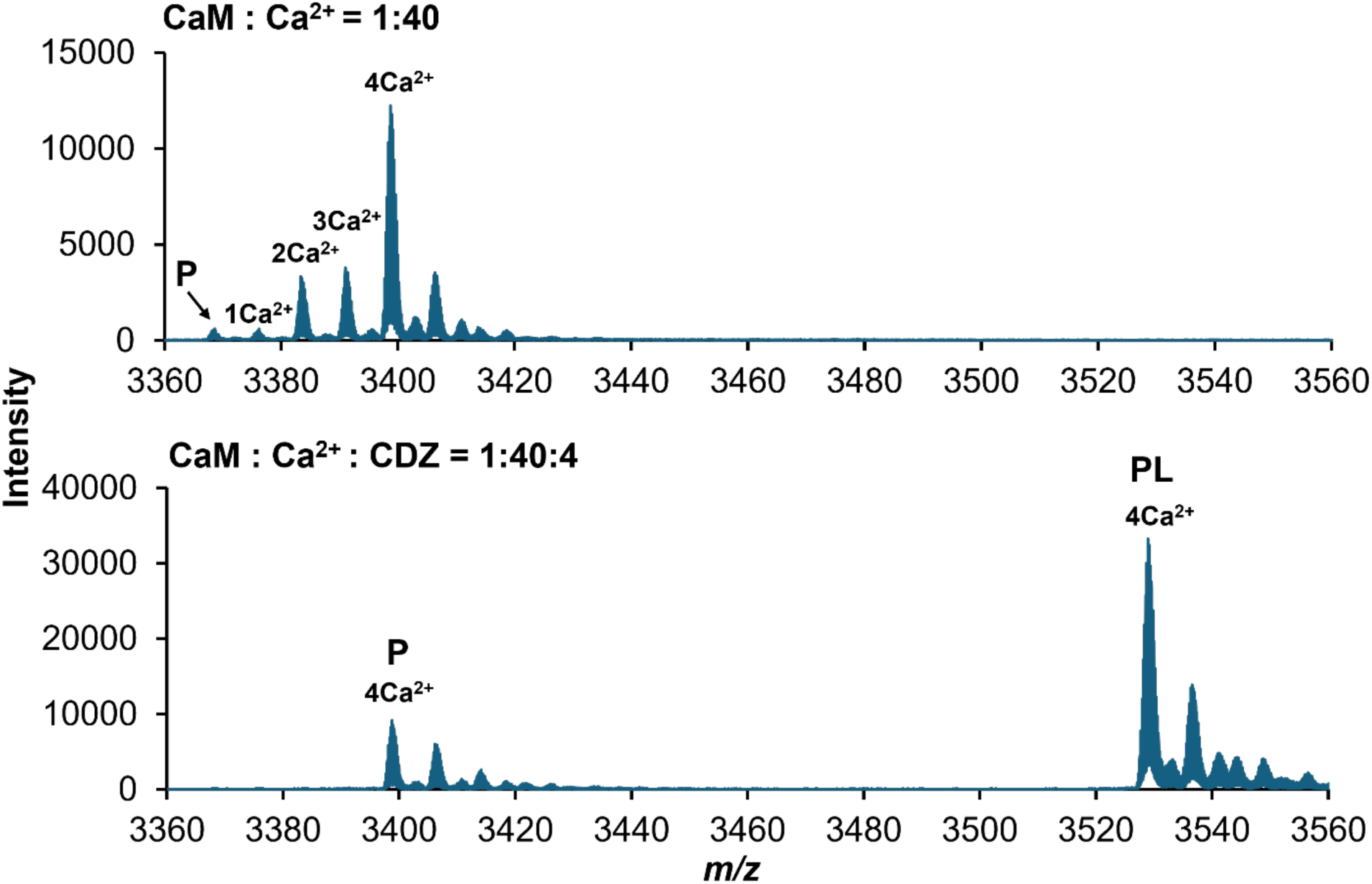
Zoomed in native mass spectra of the 6+ charge state of CaM . **Top.** CaM in the absence of CDZ (Figure 2B). **Bottom.** CaM in the presence of CDZ (Figure 2D). P and PL represent unbound and bound peaks, respectively.

Having optimized experimental conditions, the effect of CDZ binding on CaM CSD production was examined. In the mass spectrum of CaM with CDZ and Ca^2+^, the emergence of prominent, low charge state peaks in the mass spectrum correspond to the CDZ-bound CaM (Figure 2. B vs D). These results support previously published reports showing that CaM-CDZ binding is Ca^2+^-dependent.^69,92^ The Ca^2+^-triggered binding of CDZ to CaM caused a marked redistribution of the CSD peaks with a clear shift from higher to lower charge states for both bound and unbound forms. Under CDZ-free Ca^2+^ loading conditions (CaM:Ca(OAc)_2_ =1:40) as well as for apo CaM, the CSD features predominantly populated the z = 6+ to 8+ range. However, addition of CDZ and Ca^2+^ almost eliminated the 8+ population, and the 7+ and 6+ abundances were substantially reduced. Furthermore, the previously absent 5+ and 4+ ions became the dominant species in the spectra (Figure 2D).

Further analysis showed that CDZ–CaM complexes were observed as z = 6+, 5+, and 4+ species, with z = 5+ representing the dominant CDZ-bound population and z = 6+ as a minor component. Notably, the highly abundant z = 4+ charge state in the sample containing CaM, CDZ, and Ca²+ (Figure 2D) was predominantly detected as a CDZ-free species. This may seem somewhat unexpected, as z = 4+ species were absent under all other conditions. However, it is possible that the appearance of this population was also driven by non-covalent CDZ binding. Here, transient binding led to conformer stabilization that was largely preserved on the very short timescale of the ionization process. Thus, the z = 4+ population likely reflects a CDZ-induced conformational state rather than a native apo ensemble. The z = 5+ charge state also exhibited a substantial CDZ-free CaM population. To further probe the relationship between conformational state and ligand interaction, titration of CDZ with CaM in the presence of Ca²+ was performed. Analysis of the titration revealed that the fraction of bound protein increased substantially with increasing CDZ concentration for the z = 5+ species, while the z = 4+ population changed only modestly over the same concentration range (Figure S4, Table 1). This concentration dependence indicated that a fraction of the apparent CDZ-free CaM z = 5+ signal was coupled to CDZ binding rather than representing a purely ligand-free population. Because lower charge states are associated with more compact conformers in nMS, it is possible that the z = 5+ and 4+ species represent a highly compact endpoint population, whereas the z = 5+ state corresponds to a broader ensemble of binding-competent conformers that remain responsive to ligand concentration. The presence of these CDZ-free species may arise from several non-mutually exclusive mechanisms, including partial ligand loss during ionization or formation of a CDZ-induced conformational state that may transiently persist following ligand dissociation. Together, these observations demonstrate that cVSSI-based nMS can resolve ligand stoichiometry while providing insight into conformational selection and binding-associated structural stabilization.

**Table 1:** Fraction bound values for 5+ & 4+ species of CaM upon binding to CDZ at increasing ligand concentrations. I_PL_, intensity of the bound peak; I_P_, Intensity of the unbound protein peak.

| Ligand Concentration ($\mu\text{M}$) | Fraction bound $\{I_{\text{PL}}/(I_{\text{P}}+I_{\text{PL}})\}$ | |
| --- | --- | --- |
|  | 5+ | 4+ |
| 0.5 | 0.35 | 0.15 |
| 1 | 0.69 | 0.20 |
| 2 | 0.72 | 0.21 |
| 5 | 0.81 | 0.21 |
| 10 | 0.81 | 0.21 |
| 20 | 0.83 | 0.26 |

To corroborate the experimental results, a series of protein folding predictions of CaM with and without Ca^2+^ and CDZ were performed using Chai Discovery software, a deep learning structure prediction tool that can accommodate and predict protein binding with bespoke ligands.^77^ This analysis enabled the visualization of structural shifts of the protein upon ligand binding that may have been detected by cVSSI. Structural predictions showed that Ca^2+^-bound CaM remains broadly similar to the apo-CaM model, with no major conformational rearrangements. However, binding of CDZ to CaM in the presence of Ca^2+^ leads to a significant structural rearrangement, yielding a more compact and folded conformation of the protein (Figure 5). These Chai Discovery prediction results match a previously published crystal structure of CaM with CDZ (reproduced in Figure S5). This prior structural study demonstrated that binding of a single CDZ molecule to CaM can induce a large open-to-close conformational transition, yielding a compact globular complex resembling the compaction observed when holo-CaM engages with peptide targets.^69,93^ These results support the cVSSI measurements. The same study also found that CaM can bind two molecules of CDZ simultaneously. However, it was demonstrated that the second binding event had only a minimal effect on the structure and dynamics of the protein, concluding that the solution behavior is more consistent with a predominantly 1:1 stoichiometry of the interaction.^69^ Correspondingly, although our experimental data primarily found 1:1 binding, we also observed a small population of CaM binding to two CDZ molecules in the z = 6+ and 5+ species whereas no CDZ binding was observed in the z = 7+ or 8+ species under these conditions (Figure 2D). Moreover, the Chai Discovery predictions similarly showed a plausible 2:1 binding mode that matched the published crystal structure (Figure S5, Table S1). Taken together, these results demonstrate that cVSSI-based nMS provides a powerful approach to directly link ligand binding with conformational selection and structural stabilization in dynamic proteins such as CaM, consistent with results from orthogonal structure determination approaches.

**Figure 5.**
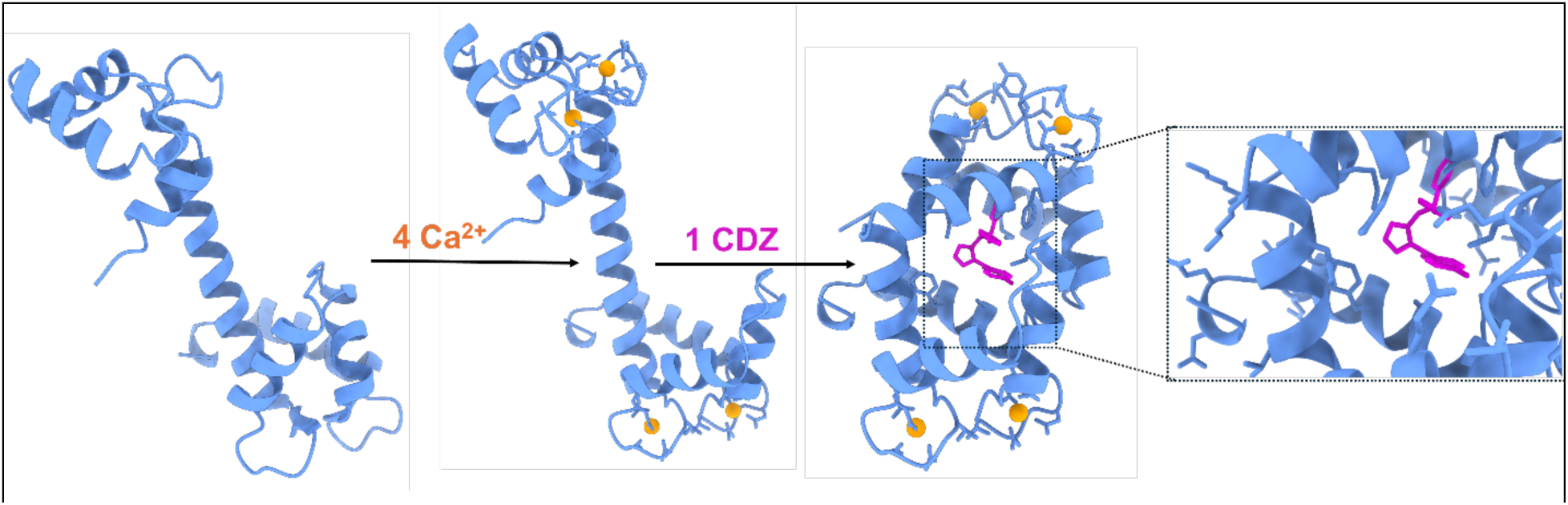
Chai Discovery predicted structural transition of CaM in different states. Left to right: Apo CaM, Ca^2+^-bound CaM, CDZ molecule-bound CaM, and zoomed-in view of the CDZ-bound region. CaM is shown in cornflower blue ribbon representation, Ca^2+^ ions are shown as orange spheres, and CDZ is shown in magenta. The models were chosen based on the highest sequence coverage to their corresponding crystal structure in the Protein Data Bank, as shown in Table S1.

### Equilibrium dissociation constant (app *K_d_*) of the interaction

In addition to characterizing dynamic conformational changes in protein structure, cVSSI can also be used to derive apparent equilibrium dissociation constants (*K_d_*) for protein-ligand interactions. In this study, the apparent *K_d_* of the CaM-CDZ interaction was determined using a ligand-based titration method. A fixed concentration of CaM (0.5 uM) and Ca^2+^ (20 uM) was titrated against increasing concentrations of CDZ, and fractions of bound protein were determined from the nMS peak-intensity ratios using Equation 1. The *K_d_* was determined using the Langmuir binding model and the quadratic binding model and the experimental values were fitted to their predicted values. Equation 2 represents the Langmuir binding model, and Figure 6A shows the non-linear binding curve where the estimated apparent *K_d_* was determined to be 261 ± 29 nM (N=3) (Table S4).

**Figure 6.**
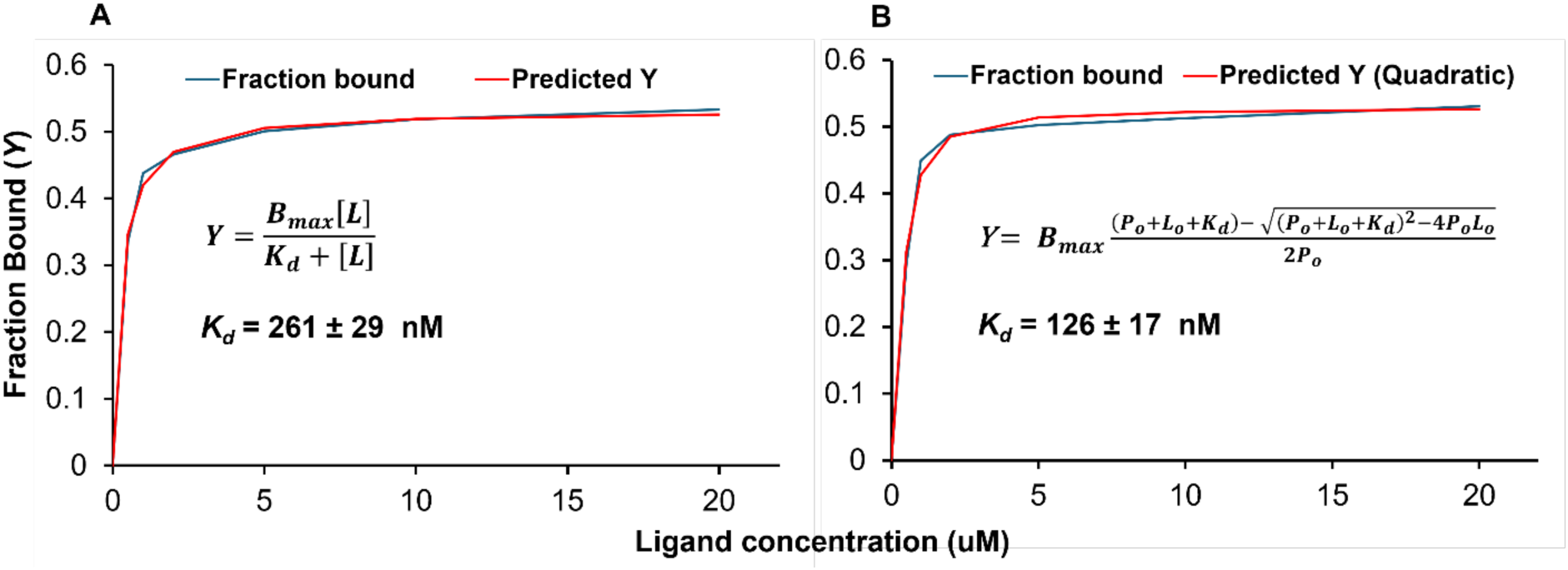
Binding curves obtained from nMS titration data. X-axis represents ligand concentrations and Y-axis the fraction of CaM bound to CDZ. Blue and red lines represent experimental fraction bound and predicted/fitted values, respectively. **A**. Data fitted with predicted values from the Langmuir binding model, yielding apparent *K_d_*: 261 ± 29 nM. **B**. Same data fitted to predicted value from the quadratic binding model, yielding apparent *K_d_*: 126 ± 17 nM. Mean ± SD from three independent experiments.

In the Langmuir binding model, the free ligand concentration was assumed to be equal to the initial concentration, which is not ideal in our system, as the free ligand concentration is reduced upon binding to the protein.^94^ To avoid ligand depletion, Equation 3 was used to determine the bound fraction in the quadratic binding model. The quadratic model is considered to account for ligand depletion under tight binding conditions.^76^ With the quadratic model, the calculated apparent *K_d_* is 126 ± 17 nM (N=3) (Figure 6B, Table S4). In both models, Microsoft Excel Solver was used to determine the best fit apparent *K_d_* where the sum of squared residuals between the predicted and experimental values are minimized (Table S2 & S3). The estimated apparent *K_d_* values from this study are within the range of previously reported CaM-CDZ affinities, which vary from nano-molar to micromolar depending on the experimental methods (e.g., ITC^69,70^, and fluorescence anisotropy^71^, and mTR-FRET assay^71^). These findings demonstrate that cVSSI-based nMS can be used to calculate apparent dissociation constants of non-covalent interactions between proteins and ligands.

### In-droplet hydrogen-deuterium exchange

The nMS data above suggest that upon binding to CDZ, CaM transforms into a more folded conformational ensemble with a population of ions having lower charge states. To further investigate the structural changes in CaM conformation driven by Ca^2+^ and CDZ binding, in-droplet HDX-MS experiments were performed. As described above, in-droplet HDX-MS offers advantages over conventional HDX-MS by enabling ultrafast labeling, thereby improving the detection of transient conformational states of dynamic proteins such as CaM. Upon exposure to D_2_O, we found the isotopic envelopes shifted towards higher *m*/*z*, reflecting incorporation of D_2_O into the exchangeable sites of the protein, as expected (Figure 7A).^48,64^ The number of incorporated D_2_O atoms was then calculated and compared between charge states as well as the unbound and CDZ-bound species within charge states to evaluate ligand-induced conformational changes and solvent accessibility (Table S5 & S6). Generally, a lower level of D_2_O incorporation is associated with reduced solvent accessibility and/or higher hydrogen bonding and can be interpreted as a more compact conformational ensemble of the protein.^95,96^ In our system, a reduction in D_2_O incorporation would also be partly attributed to a portion of the exchangeable sites of CaM being blocked by the presence of CDZ within the binding pocket.

**Figure 7.**
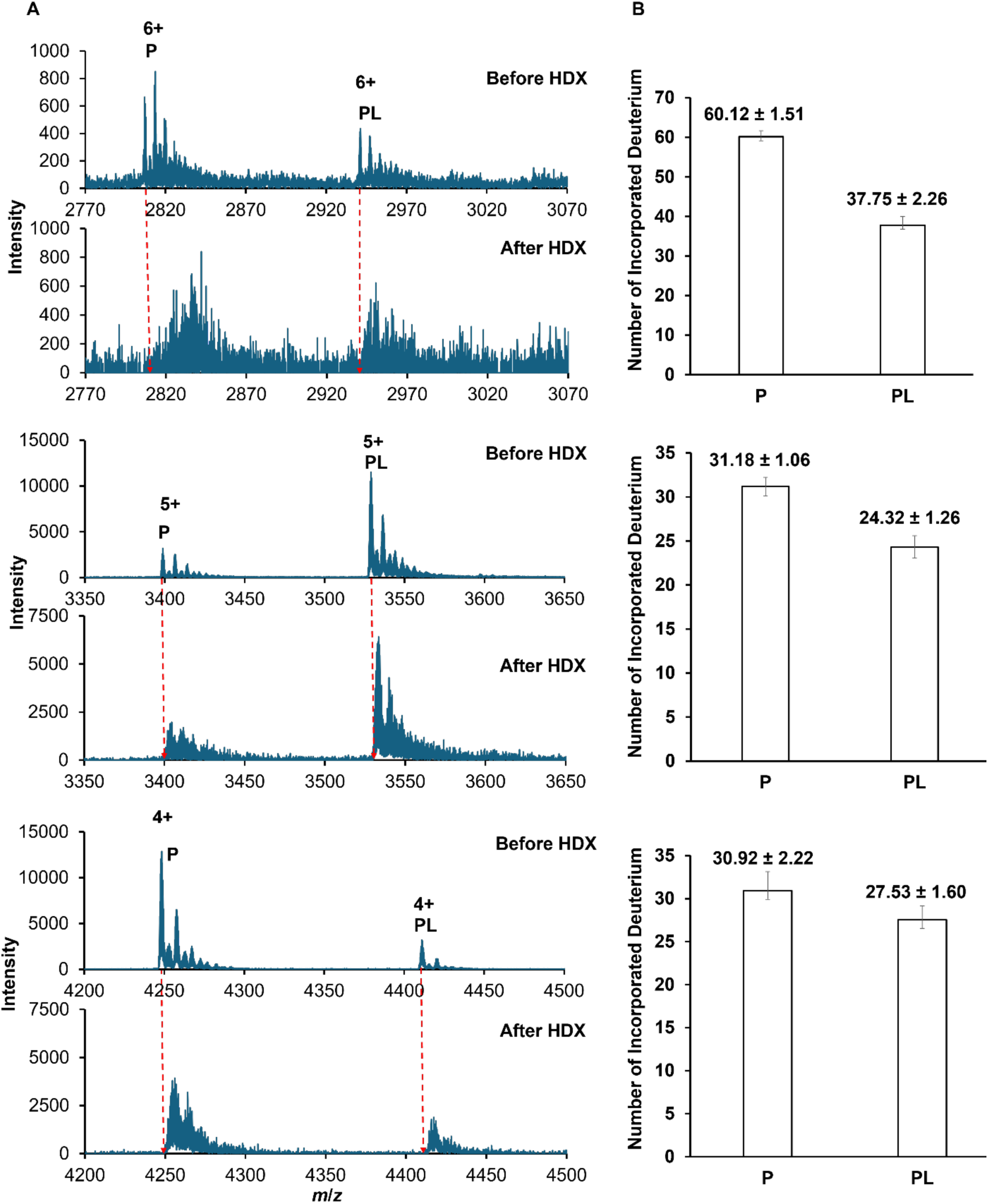
HDX-MS analysis of CaM in the absence and presence of CDZ. **A.** Zoomed-in native mass spectra of CaM before and after HDX, red dotted lines highlighting the *m*/*z* shift corresponding to deuterium incorporation. The top, middle and bottom spectra represent 6+, 5+ and 4+ charge states, respectively. **B.** Bar plot showing the average number of incorporated deuterium atoms for 6+ (top), 5+ (middle) and 4+ (bottom) species, with error bars representing standard deviation from triplicate measurements. P and PL represent unbound and CDZ-bound CaM species, respectively. Statistical significance was evaluated using paired two-tailed t-tests comparing matched P and PL replicates for each charge state. Deuterium incorporation was reduced in the CDZ-bound 6+, 5+, and 4+ CaM species with p-values of 0.00063, 0.0049, and 0.0114, respectively.

Across the entire CSD, we observed that D_2_O incorporation scaled with charge state; the 7+, 6+, 5+, and 4+ species lacking CDZ incorporated 68.5, 60.1, 31.2, and 30.9 D_2_O atoms, respectively. These findings corroborate our previous nMS results further demonstrating that both of these methodologies effectively measure stepwise conformational changes in CaM. Our in-droplet HDX-MS data suggest that the 7+ and 6+ charge states represent unfurled conformations with greater availability of D_2_O-exchangeable sites, while the 5+ and 4+ charge states represent a significantly more compact conformation. Interestingly, these data showed that the 6+, 5+ and 4+ charge states exhibited two distinct populations within each charge state representing CaM with and without a bound CDZ molecule. This phenomenon was not resolvable by nMS alone, underscoring the complementary conformational detail afforded by in-droplet HDX-MS.

When examining the two major resolved and reproducible CaM species within each charge state, representing CaM that was either bound or unbound by CDZ, we found that ligand binding consistently caused a further reduction in D_2_O incorporation. Replicate analyses within the 6+, 5+ and 4+ charge states found that CDZ binding caused a reduction in D_2_O incorporation into CaM by approximately 37%, 22% and 11%, respectively, compared to unbound CaM (Figure 7B). Because the unbound and CDZ-bound populations within each charge state share the same ionization-derived compaction state, these distinct populations are unlikely to reflect major conformational differences. Rather, the reduction in D₂O uptake upon CDZ binding within each charge state is most likely attributable to a combination of direct ligand occlusion of solvent-accessible backbone amide sites at and around the CDZ binding interface, and local structural tightening of the binding pocket induced by CDZ binding. The observed gradient in HDX protection across charge states suggests that in-droplet HDX-MS can resolve distinct stages along a continuum of ligand-induced conformational change. These data capture snapshots of CaM conformers that differ in their degree of CDZ engagement and binding interface accessibility: the more open, higher charge state conformers present a larger solvent-accessible interface to CDZ, resulting in occlusion of a greater number of exchangeable backbone amide sites upon binding, whereas the more compact lower charge state conformers offer a more constrained pocket with correspondingly fewer sites available for ligand-mediated protection. This interpretation is further supported by analysis of the 5+ charge state, where a species containing a second bound CDZ molecule was also resolved. Relative to unbound CaM, this species with two CDZ molecules showed an additional ∼10% reduction in D₂O incorporation beyond that observed for the single bound species, consistent with further ligand-mediated occlusion and structural tightening at a second binding interface (Table S5). Together, the nMS charge-state distribution and HDX-MS data for CaM·CDZ are mutually consistent with CDZ binding driving CaM toward a more compact conformation, in agreement with prior structural studies demonstrating that CDZ locks CaM into a closed state.^69^ Collectively, these results demonstrate that cVSSI-based in-droplet HDX-MS provides a sensitive and rapid means of characterizing ligand-induced conformational changes and binding interface protection in metal-dependent protein–ligand systems, revealing structural detail that complements and extends what is accessible by nMS alone.

## CONCLUSIONS

This study establishes cVSSI-based nMS, combined with in-droplet HDX-MS, as a powerful and integrative platform for resolving Ca^2+^-dependent protein–ligand interactions. Using CaM as a model system, cVSSI was found to preserve a native-like conformational ensemble. This enabled clear resolution of stepwise Ca^2+^ loading and direct observation of ligand binding. Under these experimental conditions, cVSSI produced a narrower and lower charge-state distribution for apo-CaM compared to previously reported nMS studies using conventional ESI/nESI ion sources, further supporting improved preservation of native-like structure^85,87^.

It was found that CDZ binding was strictly dependent on full Ca^2+^ occupancy, which drove a redistribution of conformational populations toward compact, low-charge states, consistent with ligand-induced structural stabilization. Following CDZ addition, partially Ca^2+^-loaded species were depleted and the fully loaded population dominated, indicating preferential binding to, and stabilization of, the Ca^2+^-saturated form of the protein. Interestingly, this binding response was not uniform across all CaM populations in the CSD. In particular, z = 5+ populations exhibited the strongest response to ligand binding with increasing CDZ concentrations, demonstrating the presence of binding-competent conformers within the ensemble. In contrast, z = 7+ and 8+ ions, which dominated the apo-CaM sample, showed little to no detectable binding. Our CDZ titration showed that the z = 7+ and 8+ peaks diminished with increasing ligand concentration, further supporting the idea that these populations represent CDZ-free CaM in extended conformations that are converted into compact species upon CDZ binding. In addition, CDZ addition generated a prominent z = 4+ population with distinct behavior: although its emergence was clearly driven by CDZ, the CDZ-free CaM species dominated this charge state. Moreover, the fraction of protein bound to CDZ was largely unaffected by increasing ligand concentration. This suggests that the z = 4+ state may arise from a CDZ-induced compact conformer that either undergoes partial ligand dissociation during ionization or corresponds to a conformational endpoint with reduced ligand accessibility and exchange.

Based on charge-state-resolved titration data, the apparent dissociation constant was estimated using Langmuir and quadratic binding models, yielding values of 261 ± 29 nM and 126 ± 17 nM, respectively, with the latter accounting for ligand depletion. The agreement between these models supports a nanomolar-affinity interaction and demonstrates that cVSSI-based nMS can provide reliable quantitative estimates of protein–ligand binding directly from native mass spectra. Complementary in-droplet HDX-MS revealed that a reduction in D₂O incorporation was correlated with lower charge states across the CSD, corroborating the nMS data and confirming a stepwise conformational compaction of CaM. Upon CDZ binding, reductions in D₂O incorporation were observed when comparing bound and unbound CaM populations within each of the 6+, 5+, and 4+ charge states, yielding protection values of approximately 37%, 22%, and 11%, respectively, consistent with ligand occlusion and local structural tightening at the binding interface. These results illustrate the capacity of in-droplet HDX-MS to simultaneously capture ligand-induced structural information across a wide range of coexisting conformational states.

Overall, these results demonstrate that the combined cVSSI-based nMS and in-droplet HDX-MS approach provides a robust framework for studying metal-dependent protein–ligand interactions. This platform enables simultaneous resolution of metal-loading states, identification of ligand binding, quantification of binding affinity, and characterization of ligand-induced conformational changes within a single experimental workflow, providing an integrated readout that is difficult to achieve with conventional nMS or HDX-MS approaches alone. As such, this approach is well-suited for interrogating structurally dynamic protein systems in which binding, metal occupancy, and conformational state are tightly coupled.

## ASSOCIATED CONTENT

### Supporting Information

Schematic of dual emitter setup for in-droplet HDX-MS. Zoomed in mass spectra of stepwise Ca^2+^ loading. Mass spectra of CDZ binding with its increased concentrations. PDB structure of CaM-Ca^2+^ and CaM-Ca^2+^-CDZ. Table of *K_d_* and HDX calculations.

## AUTHOR INFORMATION

### Corresponding Authors

**Sohag Ahmed** – Department of Chemistry, West Virginia University, Morgantown, West Virginia 26506, United States;

**Kevin C. Courtney** – Department of Molecular and Cellular Biology, University of Guelph, Guelph, ON N1G 2W1, Canada;

Former address: Department of Biochemistry and Molecular Medicine, West Virginia University, Morgantown, West Virginia 26506, United States;

Former address: Department of Chemistry, West Virginia University, Morgantown, West Virginia 26506, United States;

### Authors

**Axel Woehrling** –

**Stephen J. Valentine –**

## ACKNOWLEDGEMENTS

We are grateful to Anthony DeBastiani, Interim Manager, Bionano Research, WVU for their assistance with the mass spectrometry, and Paolo Fagone, Research Associate, Biochemistry and Molecular Medicine, WVU, for their support with protein purification and FPLC equipment. This work was supported by the National Institutes of Health (grant 1R35GM160081-01, KCC and R01GM153899, SJV and PL).

