## Supplemental for "Structural Characterization of Calcium-Dependent Calmodulin-Calmidazolium Binding using Capillary Vibrating Sharp-Edge Spray-based Native Mass Spectrometry and In-Droplet Hydrogen Deuterium Exchange Mass Spectrometry"

Supporting Information

Figure S1 2

Figure S2 3

Figure S3 4

Figure S4 5

Figure S5 6-7

Figure S6 8

Table S1 9-10

Table S2 11

Table S3 12

Table S4 13

Table S5 14

Table S6 15

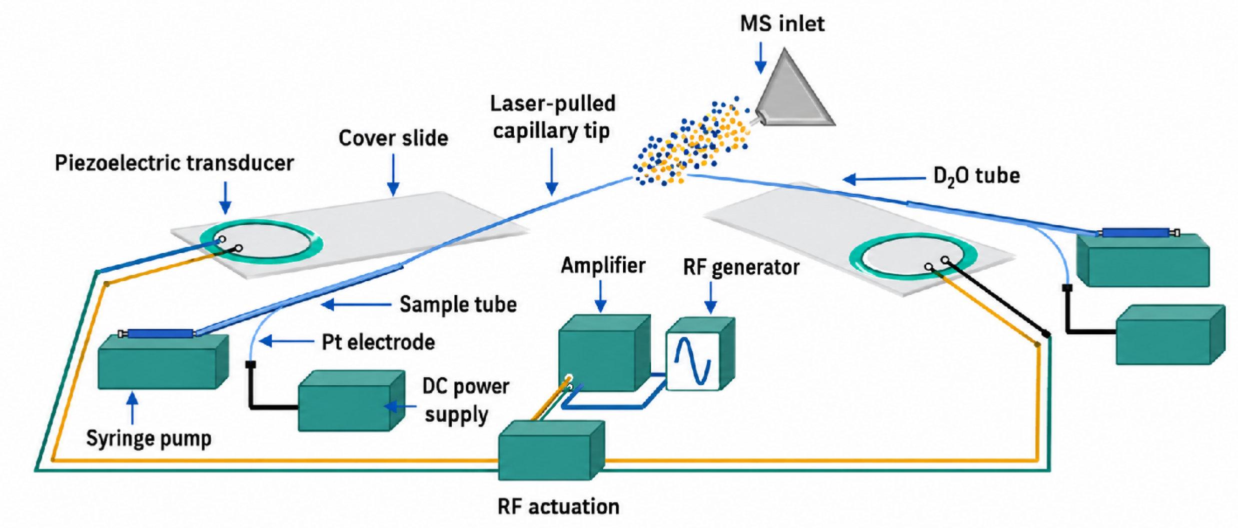


**Figure S1. Dual emitter setup containing two cVSSI devices for in-droplet HDX-MS experiments.** The left device produces droplets of the analyte solution (blue dots) and the right device produces droplets $D_2O$ (orange dots). Droplet mixing occurs in flight before entering the MS-inlet enabling rapid hydrogen-deuterium exchange. Solutions are delivered to each device using syringe pumps, and droplets are produced from vibrating laser-pulled capillary tips driven by piezoelectric transducers. An RF generator and amplifier provide the actuation signal required for capillary vibration, while a DC power supply applies voltage to the analyte solution through a platinum wire (Pt electrode) to facilitate ionization.[1,2]

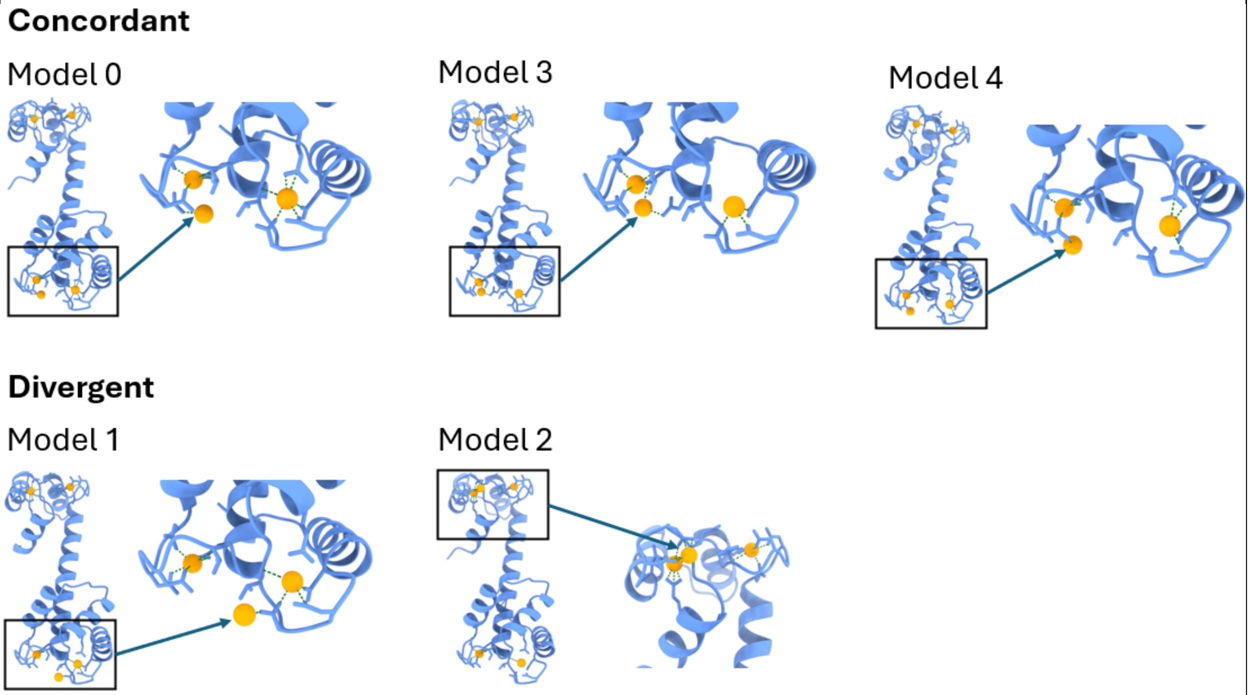


**Figure S2. Five AI-predicted CaM 5+ $Ca^{2+}$ structural models.** CaM is represented by a cornflower blue ribbon, the $Ca^{2+}$ ions are represented by orange spheres and the predicted interactions are represented by dashed green lines. Black rectangles show a zoom of the region where the fifth $Ca^{2+}$ ion binds, and green arrows point towards the fifth $Ca^{2+}$ ion. Models 0, 3, and 4 display a similar binding site for the fifth $Ca^{2+}$ ion and are regrouped as "concordant". Model 1 displays a binding site close to the one of the models 0, 3, 4 but that differs from those concordant models. Model 2 displays a completely different binding location compared to all the other models. Model 1 and model 2 are then regrouped as "divergent".

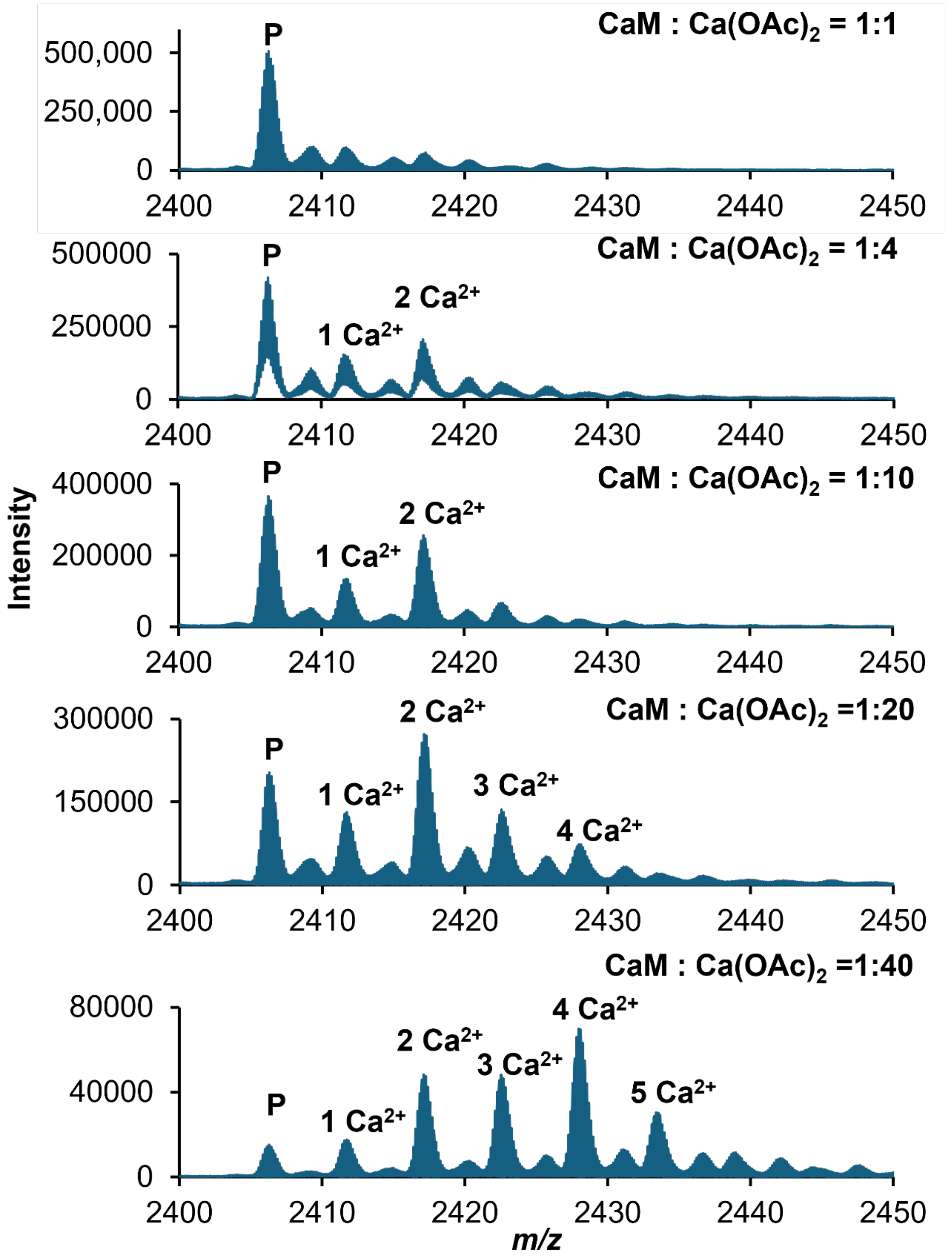


**Figure S3. Zoomed in 7+ charge state of Calmodulin at increasing concentration of $Ca(OAc)_2$.** All four calcium-binding sites of the CaM protein were occupied at a ratio of CaM (0.5 μM):$Ca(OAc)_2$ (20 μM) = 1:40.

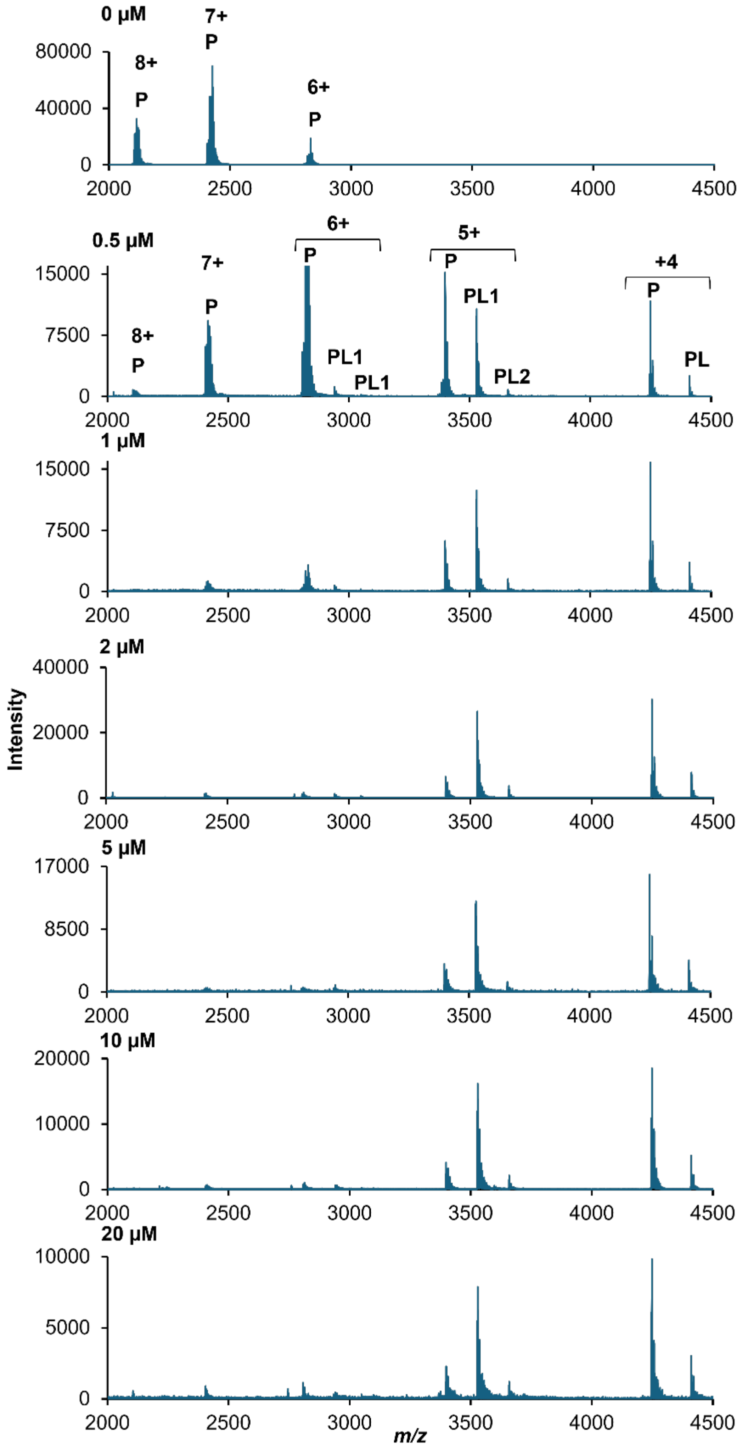


**Figure S4. Native mass spectra of CaM with increasing CDZ concentrations.** Spectra are shown from top to bottom with increasing CDZ concentration. The CaM and $Ca(OAc)_2$ concentrations were maintained at 0.5 µM and 20 µM, respectively, throughout the experiment. Peaks correspond to the observed charge states and associated protein species at each condition. P, PL1, and PL2 correspond to populations of CaM that are unbound, one and two CDZ molecules bound, respectively.

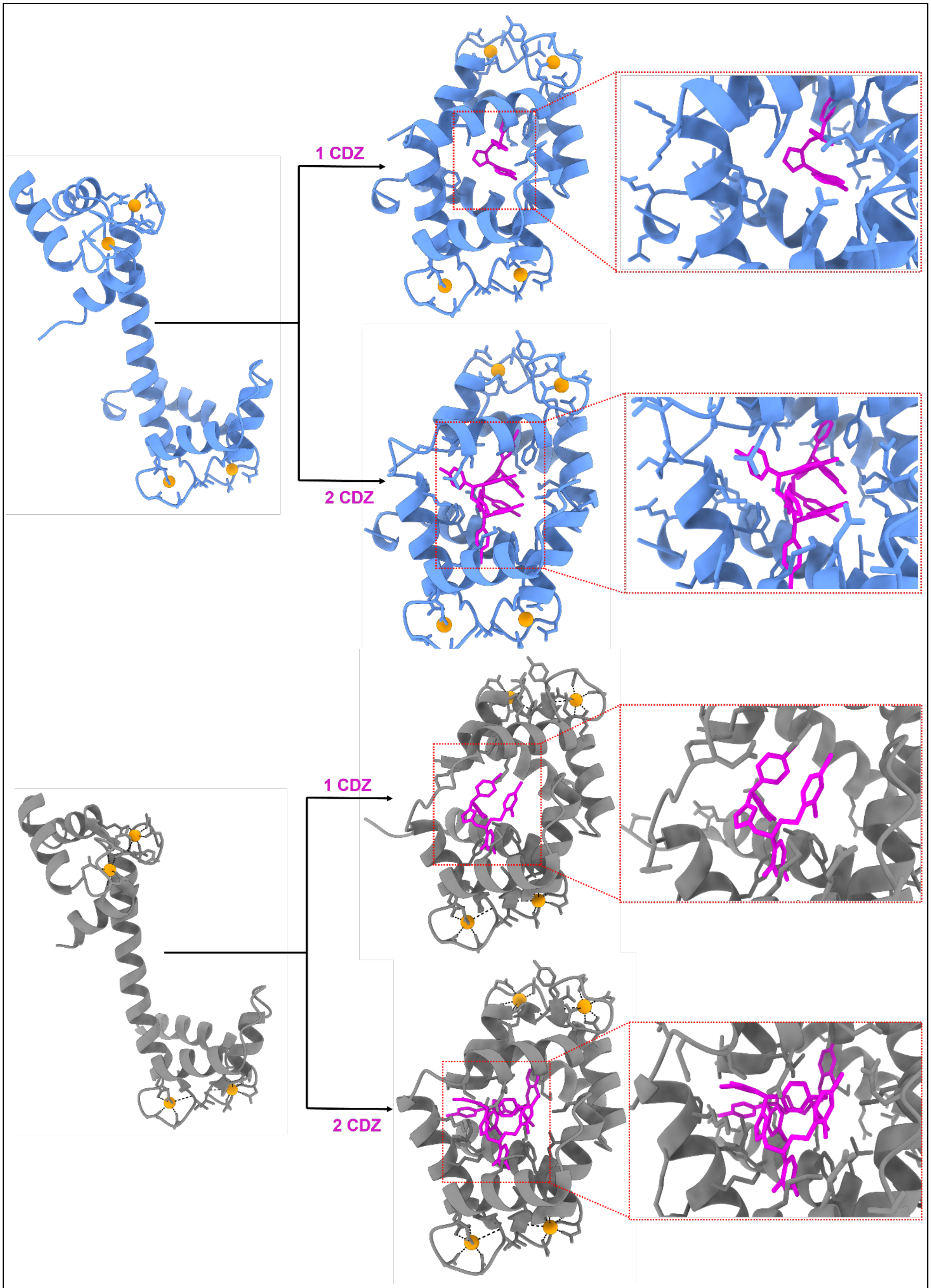


**Figure S5. Comparison between predicted and experimental crystal structures of**

**$Ca^{2+}$-bound CaM complexed with one and two CDZ molecules.** The top panel shows Chai-1 predicted structures, and the bottom panel shows structures obtained from the Protein Data Bank (PDB). In both cases, the left panel shows $Ca^{2+}$-bound, the middle panel shows one and two CDZ-bound CaM. The right panel is the enlarged view of the CDZ binding region. The CaM is shown in ribbon representation, $Ca^{2+}$-ions are represented as orange spheres, and CDZ in magenta sticks. The structures obtained from the protein data bank are $Ca^{2+}$-bound CaM (PDB ID: 1CLL[3]), one CDZ-bound CaM (PDB ID: 7PSZ[4]), and two CDZ-bound CaM (PDB ID: 7PU9[4]).

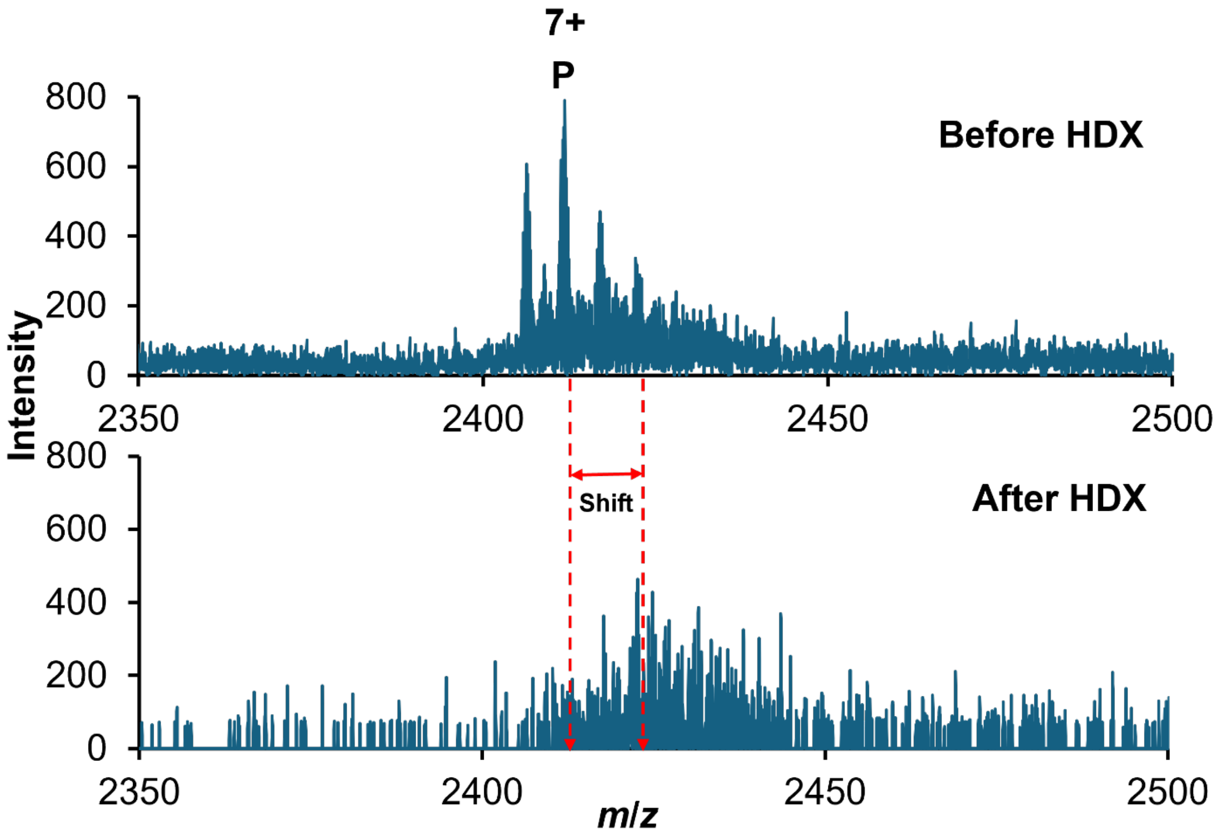


**Figure S6. Zoomed in mass spectra of 7+ CaM species before and after HDX.** The red dashed lines indicate the shift in *m*/*z* of CaM species after HDX, resulting in deuterium incorporation. P corresponds to unbound protein peaks.

**Table S1. Comparison between Chai-1 predicted structures and experimental crystal structures.** The Chai Discovery[5] generated structural models were compared to their corresponding experimental crystal structures using the MatchMaker tool in UCSF ChimeraX[6] (bb chain pairing, Needleman–Wunsch alignment algorithm, BLOSUM-62 similarity matrix, 0.3 secondary structure fraction, 18/18/6 gap open (HH/SS/other), 1 gap extend, ((6-9-6), (6-6), (4)) (HSOxHSO) secondary structure matrix, and 2 Å iteration cutoff). Chai Discovery generated five models (Model 0-4) for each of the following structures: CaM + 4 $Ca^{2+}$, CaM + 1 CDZ + 4 $Ca^{2+}$, and CaM + 2 CDZ + 4 $Ca^{2+}$. The predicted models were compared with their corresponding experimental crystal structures from the Protein Data Bank: CaM + 4 $Ca^{2+}$ (PDB ID: 1CLL[3]), CaM + 1 CDZ + 4 $Ca^{2+}$ (PDB ID: 7PSZ[4]), and CaM + 2 CDZ + 4 $Ca^{2+}$ (PDB ID: 7PU9[4]). The pruned RMSD (Å) is the root-mean-square deviation of CaM atoms positions after removal by iterative exclusion of residue pairs that are too far apart (iteration cutoff: 2 Å). The lower the pruned RMSD value, the better the structural fit of the best-aligned core. The number of pruned atom pairs corresponds to the number of atom pairs that were kept during the iterative exclusion step of the RMSD calculation. The coverage (%) highlights the fraction of CaM residues kept after pruning and is calculated as follows: Coverage=[pruned pairs / total pairs] × 100. The higher the coverage, the more structurally complete the alignment. The PDB CaM crystal structures had different residue numbers, which we considered for the calculations (CaM + 4 $Ca^{2+}$ / PDB ID 1CLL had 144 pairs, CaM + 1 CDZ + 4 $Ca^{2+}$ / PDB ID 7PSZ had 145 pairs, and CaM + 2 CDZ + 4 $Ca^{2+}$ / PDB ID 7PU9 had 144 pairs). For each structure, the model with the lowest pruned RMSD and the model with the highest coverage are highlighted to characterize structural fitness and completeness.

| Structure | Model | Pruned RMSD (Å) | Number of pruned atom pairs | Coverage (%) |
|---|---|---|---|---|
| CaM + 4 $Ca^{2+}$ | Model 0 | 1.205 | 111 | 77.1 |
| | Model 1 | **0.865** | 81 | 56.3 |
| | Model 2 | 1.031 | 130 | **90.3** |
| | Model 3 | 1.188 | 88 | 61.1 |
| | Model 4 | 1.155 | 113 | 78.5 |
| CaM + 1 CDZ + 4 $Ca^{2+}$ | Model 0 | 1.215 | 78 | 53.8 |
| | Model 1 | 1.251 | 84 | 57.9 |
| | Model 2 | 1.159 | 80 | 55.2 |
| | Model 3 | **0.606** | 54 | 37.2 |

| | | | | |
|---|---|---|---|---|
| | Model 4 | 1.263 | 92 | **63.4** |
| CaM + 2 CDZ + 4 $Ca^{2+}$ | Model 0 | **1.151** | 119 | **82.6** |
| | Model 1 | 1.192 | 105 | 72.9 |
| | Model 2 | 1.178 | 108 | 75.0 |
| | Model 3 | 1.168 | 106 | 73.6 |
| | Model 4 | 1.193 | 109 | 75.7 |

**Table S2.** Fraction of CaM bound calculated using equation (1) from the main article for two resolved and reproducible (+5 and +4) charge states across increasing CDZ concentrations. Apparent $K_d$ was determined by fitting the experimental data to the Langmuir binding model.[7] The sum of squared residuals (SSR) assesses the agreement of the fit between experimental and predicted values.

| Ligand concentration, L (μM) | Fraction bound | Predicted Y | Residual$^2$ | Fitting Parameters |
|---|---|---|---|---|
| 0 | 0 | 0 | 0 | $B_{max}$ |
| 0.5 | 0.342738829 | 0.360658711 | 0.000321122 | 0.526 |
| 1 | 0.44327044 | 0.427913197 | 0.000235845 | |
| 2 | 0.488552227 | 0.471913676 | 0.000276841 | $K_d$ (μM) |
| 5 | 0.503708699 | 0.502942956 | 5.86361E-07 | 0.229221 |
| 10 | 0.509495037 | 0.514213142 | 2.22605E-05 | |
| 20 | 0.526183644 | 0.520039791 | 3.77469E-05 | |
| | | SSR | 0.000894402 | |

**Table S3.** Fraction of CaM bound calculated using equation (1) from the main article for two resolved and reproducible (+5 and +4) charge states across increasing CDZ concentrations. Apparent Kd was obtained by fitting the binding data to the Quadratic binding model.[8] The sum of squared residuals (SSR) assesses the agreement of the fit between experimental and predicted values.

| **Ligand concentration, $L_0$ (μM)** | **Fraction bound** | Predicted Y (Quadratic) | Residual | $Residual^2$ | Fitting Parameters |
|---|---|---|---|---|---|
| 0 | 0 | 0 | 0 | 0 | |
| 0.5 | 0.342739 | 0.334043 | 0.008695786 | 7.56167E-05 | Protein Conc.($P_o$) |
| 1 | 0.443270 | 0.446378 | -0.003107523 | 9.6567E-06 | 0.5 μM |
| 2 | 0.488552 | 0.494985 | -0.006432812 | 4.13811E-05 | $B_{max}$ |
| 5 | 0.503709 | 0.51756 | -0.01385103 | 0.000191851 | 0.53 |
| 10 | 0.509495 | 0.524022 | -0.014526619 | 0.000211023 | |
| 20 | 0.526184 | 0.527069 | -0.000885567 | 7.84229E-07 | $K_d$ (μM) |
| | | | SSR | 0.000530312 | 0.108446 |

**Table S4.** Apparent $K_d$ values obtained from triplicate CaM-CDZ native MS titration experiments using Langmuir and quadratic binding models.

| Experiment | Langmuir $K_d$ (nM) | Quadratic $K_d$ (nM) |
|---|---|---|
| Replicate 1 | 284.52 | 142.54 |
| Replicate 2 | 229.22 | 108.45 |
| Replicate 3 | 269.67 | 125.72 |
| **Mean ± SD** | **261.14 ± 28.62** | **125.57 ± 17.05** |

**Table S5.** Deuterium incorporation calculated from HDX envelope center (*m*/*z)* shifts for unbound and CDZ-bound CaM at the 7+ to 4+ charge states across triplicate experiments. The number of incorporated deuterium was calculated for two binding sites observed in the 5+ species, where the differences in the number of incorporated deuterium were calculated relative to 1st CDZ-bound population and the percentage was calculated comparing unbound CaM population. For the 7+ charge state, the number of incorporated deuterium was calculated for the unbound protein population only.

| Experiments | Condition | After HDX m/z | Before HDX *m/z* | Δ *m/z* | Charge states | Incorporated deuteriums (D) | Reduction (D) | Reduction (%) |
|---|---|---|---|---|---|---|---|---|
| 1 | Unbound | 4256.78 | 4248.2 | 8.37 | 4 | 33.46 | 4.12 | 12.31 |
| | Bound | 4418.65 | 4411.31 | 7.34 | | 29.34 | | |
| 2 | Unbound | 4255.51 | 4248.2 | 7.33 | | 29.32 | 3.02 | 10.29 |
| | Bound | 4417.89 | 4411.31 | 6.58 | | 26.31 | | |
| 3 | Unbound | 4255.91 | 4248.2 | 7.49 | | 29.97 | 3.03 | 10.11 |
| | Bound | 4418.04 | 4411.31 | 6.73 | | 26.94 | | |
| 1 | Unbound | 3405.15 | 3398.92 | 6.23 | 5 | 31.16 | 7.81 | 25.07 |
| | Bound | 3533.87 | 3529.18 | 4.68 | | 23.35 | | |
| 2 | Unbound | 3405.37 | 3398.92 | 6.45 | | 32.25 | 6.51 | 20.2 |
| | Bound | 3534.35 | 3529.18 | 5.16 | | 25.74 | | |
| 3 | Unbound | 3404.95 | 3398.92 | 6.02 | | 30.12 | 6.24 | 20.73 |
| | Bound | 3533.97 | 3529.18 | 4.79 | | 23.88 | | |
| 1 | Bound | 3663.51 | 3659.48 | 4.03 | 5 | 20.14 | 3.21 | 10.30 |
| 2 | Bound | 3663.98 | 3659.48 | 4.49 | | 22.47 | 3.27 | 10.14 |
| 3 | Bound | 3663.61 | 3659.48 | 4.12 | | 20.61 | 3.27 | 10.86 |
| 1 | Unbound | 2835.56 | 2825.78 | 9.79 | 6 | 58.71 | 23.46 | 39.96 |
| | Bound | 2956.04 | 2950.16 | 5.87 | | 35.25 | | |
| 2 | Unbound | 2836.06 | 2825.78 | 10.28 | | 61.71 | 22.07 | 35.76 |
| | Bound | 2956.77 | 2950.16 | 6.61 | | 39.64 | | |
| 3 | Unbound | 2835.77 | 2825.78 | 9.99 | | 59.96 | 21.58 | 36.00 |
| | Bound | 2956.56 | 2950.16 | 6.39 | | 38.37 | | |
| 1 | Unbound | 2428.91 | 2418.84 | 10.07 | 7 | 70.51 | No drug-bound species. | |
| 2 | Unbound | 2428.50 | 2418.84 | 9.66 | | 67.64 | | |
| 3 | Unbound | 2428.46 | 2418.84 | 9.62 | | 67.37 | | |

**Table S6. Quantitative summary of deuterium uptake for unbound and CDZ-bound CaM at the 6+, 5+, and 4+ charge states, with unbound 7+ CaM population included for comparison.** Values were calculated from HDX envelope center m/z shifts across triplicate experiments. The difference and reduction percentages were calculated relative to the corresponding unbound CaM value for each replicate, except for the 2nd CDZ-bound population. For the 2nd bound population, the reduction percentage was calculated relative to the unbound and the difference was calculated relative to the 1st CDZ-bound population.

| Charge state | | Experiment | Deuterium uptake before CDZ binding (D) | Deuterium uptake after CDZ binding (D) | Difference (D) | Reduction (%) |
|---|---|---|---|---|---|---|
| **4+** | | 1 | 33.46 | 29.34 | 4.12 | 12.31 |
| | | 2 | 29.32 | 26.31 | 3.02 | 10.29 |
| | | 3 | 29.97 | 26.94 | 3.03 | 10.11 |
| | | **Mean ± SD** | **30.92 ± 2.22** | **27.53 ± 1.60** | **3.39 ± 0.63** | **10.90 ± 1.22** |
| **5+** | **1st** | 1 | 31.16 | 23.35 | 7.81 | 25.07 |
| | | 2 | 32.25 | 25.74 | 6.51 | 20.20 |
| | | 3 | 30.12 | 23.88 | 6.24 | 20.73 |
| | | **Mean ± SD** | **31.18 ± 1.06** | **24.32 ± 1.26** | **6.86 ± 0.84** | **22.00 ± 2.67** |
| | **2nd** | 1 | 23.35 | 20.14 | 3.21 | 10.30 |
| | | 2 | 25.74 | 22.47 | 3.27 | 10.14 |
| | | 3 | 23.88 | 20.61 | 3.27 | 10.86 |
| | | **Mean ± SD** | **24.32 ± 1.26** | **21.07 ± 1.23** | **3.25 ± 0.03** | **10.43 ± 0.38** |
| **6+** | | 1 | 58.71 | 35.25 | 23.46 | 39.96 |
| | | 2 | 61.70 | 39.63 | 22.07 | 35.76 |
| | | 3 | 59.95 | 38.37 | 21.59 | 36.00 |
| | | **Mean ± SD** | **60.12 ± 1.51** | **37.75 ± 2.26** | **22.37 ± 0.97** | **37.24 ± 2.35** |
| **7+** | | 1 | 70.51 | **Not Applicable** | | |
| | | 2 | 67.63 | | | |
| | | 3 | 67.37 | | | |
| | | **Mean ± SD** | **68.51 ± 1.73** | | | |

(2) DeBastiani, A.; Majuta, S. N.; Sharif, D.; Attanayake, K.; Li, C.; Li, P.; Valentine, S. J. Characterizing Multidevice Capillary Vibrating Sharp-Edge Spray Ionization for *In-Droplet*

Hydrogen/Deuterium Exchange to Enhance Compound Identification. *ACS Omega* **2021**, *6* (28), 18370–18382. https://doi.org/10.1021/acsomega.1c02362.
(3) Chattopadhyaya, R.; Meador, W. E.; Means, A. R.; Quiocho, F. A. Calmodulin Structure Refined at 1.7 Å Resolution. *Journal of Molecular Biology* **1992**, *228* (4), 1177–1192. https://doi.org/10.1016/0022-2836(92)90324-D.
(4) Léger, C.; Pitard, I.; Sadi, M.; Carvalho, N.; Brier, S.; Mechaly, A.; Raoux-Barbot, D.; Davi, M.; Hoos, S.; Weber, P.; Vachette, P.; Durand, D.; Haouz, A.; Guijarro, J. I.; Ladant, D.; Chenal, A. Dynamics and Structural Changes of Calmodulin upon Interaction with the Antagonist Calmidazolium. *BMC Biol* **2022**, *20* (1), 176. https://doi.org/10.1186/s12915-022-01381-5.
(5) Chai Discovery; Boitreaud, J.; Dent, J.; McPartlon, M.; Meier, J.; Reis, V.; Rogozhnikov, A.; Wu, K. Chai-1: Decoding the Molecular Interactions of Life. Synthetic Biology October 11, 2024. https://doi.org/10.1101/2024.10.10.615955.
(6) Goddard, T. D.; Huang, C. C.; Meng, E. C.; Pettersen, E. F.; Couch, G. S.; Morris, J. H.; Ferrin, T. E. UCSF ChimeraX: Meeting Modern Challenges in Visualization and Analysis. *Protein Science* **2018**, *27* (1), 14–25. https://doi.org/10.1002/pro.3235.
(7) Li, J.; Ma, X.; Wang, X.; Hu, X.; Fang, S.; Jin, G.; Liu, K.; Dong, Z. Mutations Found in Cancer Patients Compromise DNA Binding of the Winged Helix Protein STK19. *Sci Rep* **2024**, *14* (1), 14098. https://doi.org/10.1038/s41598-024-64840-9.
(8) Liu, W. H.; Zheng, J.; Feldman, J. L.; Klein, M. A.; Kuznetsov, V. I.; Peterson, C. L.; Griffin, P. R.; Denu, J. M. Multivalent Interactions Drive Nucleosome Binding and Efficient Chromatin Deacetylation by SIRT6. *Nat Commun* **2020**, *11* (1), 5244. https://doi.org/10.1038/s41467-020-19018-y.
